# A Toolbox for the Personalization of Plasmonic Photothermal Therapy in Orthotopic Mouse Tumor Models

**DOI:** 10.1101/2025.05.02.651947

**Authors:** Pablo R. Fernández-Esteberena, Clara Vilches, Jordi Morales-Dalmau, Miguel Mireles, María del Mar Martínez-Lozano, Ignacio de Miguel, Oriol Casanovas, Romain Quidant, Turgut Durduran

## Abstract

The clinical translation of plasmonic photothermal therapy is hindered by a lack of reliable personalization protocols. In this study we implemented a non-invasive toolbox to assess how tumor optical and hemodynamic properties can serve as markers of progression and be used in personalized simulations to extrapolate therapy conditions, both necessary steps for individualized treatment planning. The toolbox integrated near-infrared diffuse optical monitoring techniques, thermal imaging, *ex vivo* assays and physical simulations. It was applied to patient-derived orthotopic clear cell renal cell carcinoma mouse models, at 1 cm^3^ volumes, providing clinically more relevant conditions than those of commonly used models. Gold nanorods (GNRs) with absorption tuned to the optical window and fixed irradiation conditions were employed. Therapy safety and efficacy were established using standard methods and pre-GNR injection, pre-therapy and post-therapy measurements obtained with the toolbox were analyzed to model therapy progression. Results revealed an association between tumor water concentration and accumulated GNR fraction in the tumor, a correlation between the pre-therapy tumor absorption coefficient and the maximum skin temperature reached, and a link between simulated treated volume estimates and progression-free survival, among other findings. These results demonstrate the capabilities of optical measurements to model the outcome and explain the mechanisms involved in the therapy, advancing towards a personalization strategy.

## 1 Introduction

Nanoparticle (NP) applications in the field of oncology have been studied for decades, trying to exploit their capacity for localized and targeted effects with the aim of improving the screening and treatment of cancer [1–3]. Yet, their translation into the clinical management of cancer has been slow for multiple reasons, including challenges in NP delivery to the tumor, their potential for toxicity, the complexity of their clearance routes, the unknown interactions with the biological environment and – a central problem in the field – the intrinsic heterogeneity of malignant tumors.

Plasmonic photothermal therapy (PPTT) is one of such implementations, aimed to treat tumors or work in synergy with other therapies. It makes use of the high efficiency of biocompatible metallic NPs (mainly gold) to absorb light and act as localized sources of heat [4–7]. The NPs are either injected into tumor tissue or into the bloodstream, to later accumulate in the tumor due to its defective vasculature and lymphatic system [8, 9]. The external illumination of the tissue then excites the collective oscillations of the surface electrons in the NPs, phenomenon known as surface plasmon resonances, resulting in the enhanced heating of the tissue, which in turn can provoke physiological changes, induce cell death and reduce tumor volume and/or growth rate. As a localized ablation method it is a potential treatment for unresectable tumors that are resistant to chemotherapy and radiation therapy (for example in the liver, kidneys, bones, lungs, esophagus or rectum) [10], whereas in a mild hyperthermia regime, it can be used neo-adjuvantly to sensitize cells to ionizing radiation therapy [11], enhance the uptake of anticancer drugs [12], improve photosensitizer delivery and oxygen supply for photodynamic therapy (PDT) [13, 14] or reduce tumor margins for surgery [15].

Numerous preclinical studies have proven the efficacy of PPTT ablation *in vivo* on tumor-bearing mice with diverse gold NP shapes, sizes, surface functionalizations and tumor types (reviewed in Ref. [16]). Similar findings were reported in larger animal models with spontaneous [17–20] and inoculated [21] tumors. These preclinical studies have generally shown the elimination of the tumor burden while reporting no toxic effects derived from the use of nanomaterials. However, the vast majority (see [16]) consisted in (a) subcutaneous tumors, lacking the complex microenvironment and interconnected network of vessels that would grow in the primary host organ [22], and, (b) tumors smaller than the characteristics diffusion lengths of light and heat (*<*6 mm in diameter) [23, 24]. These tumors represent, due to this over-simplification, only the first steps towards demonstrating the potential of PPTT for clinical application. It is also important to note that there are no standardized protocols for irradiation that try to account for the large number of therapy parameters involved, including the nanoparticle dosage, the illumination method, the illumination power and the exposure time.

Moreover, data from PPTT in humans is scarce. To date, three clinical trials have been approved using silica-gold nanoshells to treat prostate tumors, head and neck cancers and metastatic lung cancer (NCT04240639, NCT00848042, NCT01679470, respectively). The only published group study, by Rastinehad *et al*. [25], demonstrated the feasibility and the safety of the approach with data from fifteen prostate cancer patients. In addition, Kadria-Vili *et al*. published a prostate cancer PPTT case study with one year follow-up with good results [26]. Nonetheless, its wider applicability and long-term efficacy are yet to be demonstrated.

These facts motivated the development and implementation of a comprehensive toolbox, akin to those used along other cancer therapies, with the aim of informing the personalization of PPTT while addressing the gap between experiments and clinical translation. This meant it must be capable of obtaining relevant information about the tumor to treat and relating it to the critical aspects of the therapy: NP delivery, hyperthermia and therapy outcome.

In the literature, imaging approaches like X-ray computed tomography (CT), magnetic resonance imaging (MRI) or photoacoustic imaging (PAI) have been proposed to measure NP accumulation or tissue temperature to guide or tailor the therapy and accelerate its progression to the clinics [27–29]. However, these are not available in all clinics and despite their complexity and cost, provide incomplete information to understand the large variability in the outcome of the therapy.

In this sense, diffuse optical techniques offer advantages for therapy personalization. They provide innocuous, non-invasive means to probe centimeters deep into the tissue with NIR light in the optical window (approximately 650 – 950 nm), where light absorption by tissues is typically low, and extract information about their structure, optical properties and hemodynamics. They do not require contrast agents, are compact and low-cost in comparison to more spread imaging techniques [23, 30]. They have been broadly applied to the study of cancer therapy, mainly chemotherapy and PDT in humans [31–33] and preclinical models [34–41], but, to the best of our knowledge, have not been used to investigate the dynamics of PPTT. In the application of this therapy, the scattering and absorption properties of tumors are crucial because they determine the transport of treatment light and the generation of heat in the tissue, while tumor hemodynamics have been linked to tumor evolution and NP delivery in some models [9, 42–44]. In this direction, He, Li *et al*. have implemented diffuse optical tomography with an MRI-compatible transrectal probe to monitor tissue hemodynamics during photothermal therapy [45] and demonstrated its use in dog prostate (without NPs or a tumor) [46].

In this work, we have developed a toolbox comprised of near-infrared (NIR) diffuse optical monitoring techniques, infrared temperature monitoring, *ex vivo* assays and Monte Carlo light transport and heat diffusion simulations, adapted to *in vivo* mouse experiments of PPTT on large tumors (∼1 cm). In particular, the optical devices combines broadband diffuse reflectance spectroscopy (DRS) and diffuse correlation spectroscopy (DCS) [47, 48]. DRS can recover oxy- and deoxy-hemoglobin, water and/or lipid concentrations by deriving them from the absorption and scattering properties of the tissue. DCS uses fluctuations in the intensity of coherent laser light that has traveled through the tissue to estimate the motion of red blood cells, which indicates blood flow.

The development and testing of this toolbox was carried out on extensive PPTT experiments using a fixed set of therapy conditions on a patient-derived orthotopic clear cell renal cell carcinoma (ccRCC) mouse model, treated at a volume of 1 cm^3^, providing a more realistic scenario than the small subcutaneous tumors utilized in the literature. This model has been extensively used for the study of antiangiogenic therapy [49], including with diffuse optical longitudinal monitoring [47]. Moreover, we used gold nanorods (GNR) coated with polyethylene glycol (PEG) with a plasmon resonance tuned to the optical window (∼ 810 nm), allowing on one side the plasmonic excitation deeper below the skin, not limiting the generation of heat to the most superficial layer [23] and, on the other, the quantification with DRS of the GNR concentration that accumulated in tumor tissue, as we have shown is possible in [48].

We hypothesized that optical measurements along the different steps of the PPTT protocol are relevant markers for therapy and allow to account for some of the heterogeneity in outcome. We have thus related tumor properties measured before GNR injection to the GNR concentration delivered to the tumor, and the properties measured pre- and post-treatment to the temperatures reached on the tumor with irradiation and the outcome variables of mouse survival and progression-free survival (PFS). Furthermore, the optical properties of probed tumors were used as inputs for the simulation of light and heat transport, which was validated against our experimental results and then used to extrapolate the treatment conditions to other powers, irradiation times and absorption coefficient values. In doing so, we have demonstrated how the different techniques in the toolbox contribute to obtaining a set of rules to adjust PPTT with the outlook of optimizing its efficacy.

## 2 Methods

### 2.1 Synthesis and functionalization of gold nanoparticles

All GNR suspensions were manufactured in-house using a seed-mediated method as reported in [50], based on methods by references [51, 52]. GNRs of 11 *×* 44 nm in average (size dispersion of 15%) were grown in cetyltrimethylammonium bromide (CTAB, Sigma-Aldrich) suspension and coated with alpha-thio-omega-carboxy polyethylene glycol (HS-PEG-COOH, Iris Biotech GmbH) at 1 mg/ml, obtaining pegylated GNR (GNR–PEG).

Optical absorption spectra of GNR–PEG suspensions were measured before injection for each batch, showing a surface plasmon resonances around 810 nm.

Further details of GNR synthesis in CTAB, surface modifications and optical measurements are described in Section 1 of the Supporting information.

### 2.2 Orthotopic tumor model

The animal procedures were carried out at the animal facility of the Institut de Recerca Biomèdica de Bellvitge (IDIBELL, L’Hospitalet de Llobregat, Barcelona, Register number B9900010), which fulfilled the government regulations on the use of experimentation animals.

All protocols were reviewed and approved by IDIBELL’s Animal Experimentation Ethics Committee and by the corresponding Department of Generalitat de Catalunya (DAAM#4899).

In a similar way as described in reference [48], patient-derived renal cell carcinomas were initially grown in a cohort of athymic nude mice (immunodeficient, 4-6 weeks, Harlan Laboratories, Spain) by injecting 10^6^ cells of 786-O cell line (ATCC CRL-1932) directly into their left renal cortex. When the tumor reached 2 cm^3^, the host mouse was sacrificed, the tumor was excised, cut in small pieces and stored for implantation on new mice.

For subsequent mice, the left kidney was exposed, the piece sutured with polypropylene 7-0 surgical colorless suture (BBraun Surgical, Spain) and the incision closed with surgical staples or sutures. All procedures were done under inhaled anesthesia using 2 l/min oxygen flow with 4% isoflurane for induction and 1.5 l/min oxygen flow with 2% isoflurane for maintenance.

Mice were housed in a 12 h light-dark cycle, with controlled temperature and humidity, and free access to food and water.

When the tumor reached 1 cm^3^ (around two weeks after implantation), mice were randomly separated in groups according to the corresponding experimental protocol.

### 2.3 Evaluation of GNR biodistribution

A single intravenous dose of GNR–PEG (21 nM, 52 *·* 10^6^ g/mol molar mass, corresponding to a total quantity of 109 µg of Au) was given to athymic nude tumor-bearing mice (2 months-old). Twenty-four hours after the administration of GNR–PEG, animals were deeply anesthetized (5% isoflurane, 1,5 l/m oxygen flow) and 1 ml of blood was collected via cardiac puncture.

Tumor, liver, spleen, heart, lung and blood samples were collected and kept at -80°C for gold content analysis. Samples were diluted with an aqueous solution of nitric acid (HNO_3_) 2% weight*/*weight and analyzed for gold concentration by inductively coupled plasma mass spectrometry (ICP-MS; Agilent 7500, Agilent Technologies, USA). Gold quantity determination was done at Unitat d’Anàlisi de Metalls, Centres Cientifics i Tecnològics de la Universitat de Barcelona (CCIT-UB, Barcelona).

### 2.4 Chronic toxicity study

Chronic toxicity of GNR–PEG injection was assessed in CD-1 mice, commonly used in toxicology for safety and efficacy testing. CD-1 adult mice (Crl:CD1, 6-9 weeks old; Charles River Laboratories, USA) were injected with a single, intravenous dose of 100 µl of 21 nM GNR–PEG (*n* = 8), or equal volume of phosphate-buffered saline (PBS) for the control group (*n* = 6). Body weight and general welfare of animals was assessed twice a week. Six months after GNR–PEG or PBS administration, intracardiac puncture was performed under deep inhaled isoflurane anesthesia to obtain up to 1 ml of blood, collected in EDTA tubes for hematology analysis. Plasma for biochemistry analysis was obtained by collecting blood in lithium heparin tubes, centrifuged at 4°C, 3500 rpm during 10 min. Blood analyses were performed in an hematology analyzer (ADVIA 120 hematology instrument, Siemens Healthineers, USA; a detailed list of analytes can be found in Section 2 of the Supporting information. At the time of necropsy, macroscopic evaluation was done prior to the harvesting of organs, which were kept in paraformaldehyde for further histological analysis. Necropsy, sample collection, histopathological examination and analyses were performed by the Integrated Services of Laboratory Animals (SIAL) at Universitat Autònoma de Barcelona (UAB, Bellaterra, Barcelona).

### 2.5 Plasmonic photothermal therapy protocol

The timeline of PPTT experiments is summarized in Figure **1A**. Orthotopic xenograft renal tumors were implanted in nude mice and allowed to grow as described in section 2.2. When the tumors reached 1 cm^3^ in volume, the mice were randomly divided into three groups: (a) those receiving an intravenous injection of 21 nM GNR–PEG and therapy laser irradiation (GNR– PEG group), (b) those receiving an intravenous injection of PBS and therapy laser irradiation (PBS group) and (c) those receiving no injection nor irradiation (control group). The intravenous injections of both PBS and GNR–PEG were performed via tail vein, using 100 µl of injected volume. The injected dose of GNR–PEG corresponded to 109 µg of Au.

Twenty-four hours post-administration, animals were anesthetized with isoflurane (4% induction dose with 2 l/min oxygen flow, and 2% maintenance dose with 1.5 l/min oxygen flow) and moved to a thermal bed inside an enclosed cabinet, shown in **Figure 1B**.

**Figure 1:**
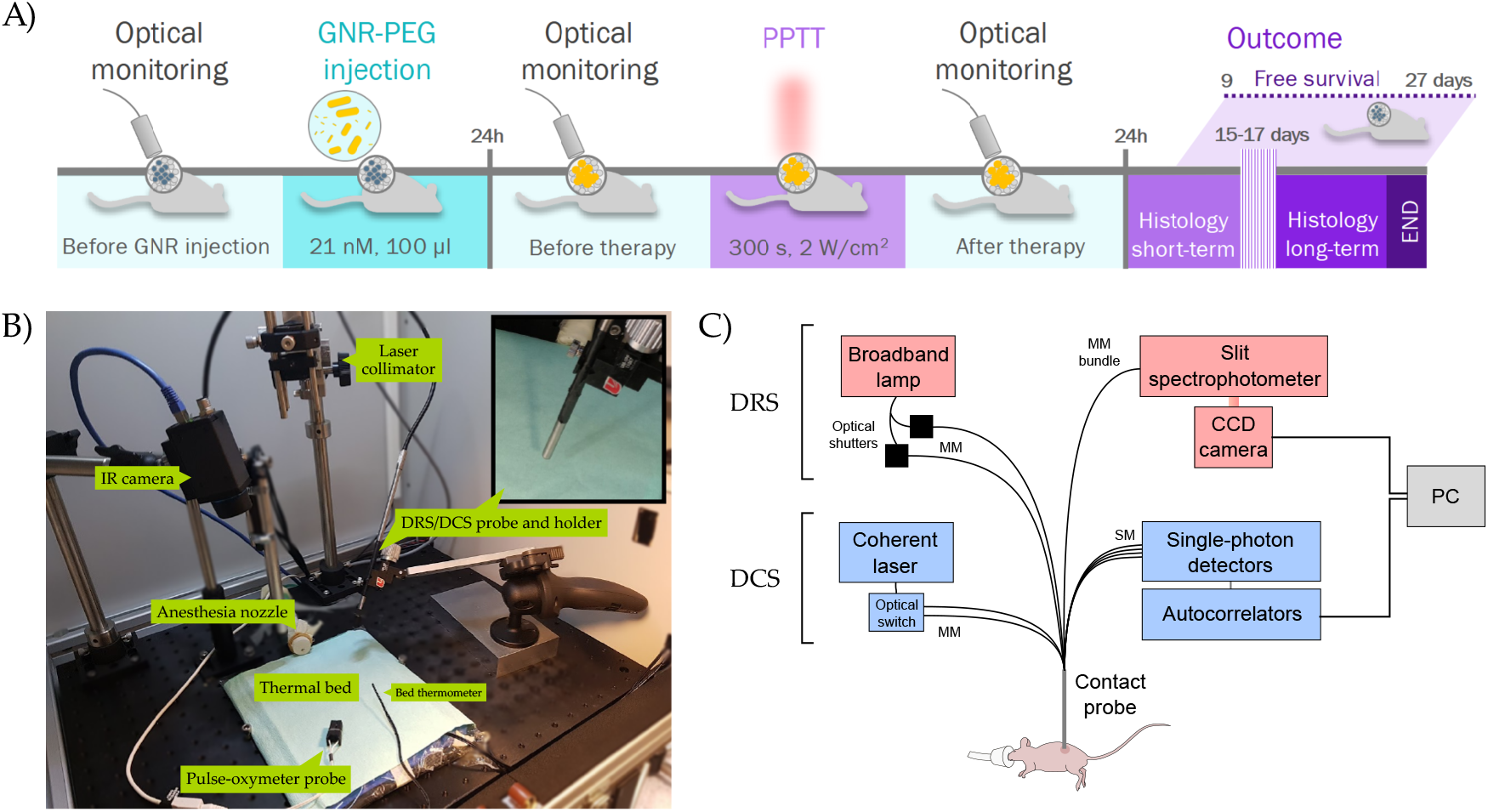
Summary of experimental protocol and setups. (A) Timeline for PPTT experiments and therapy parameters used. (B) Picture of the inside of cabinet used for PPTT experiments and components. (C) Schematic of the hybrid DRS/DCS system. CCD: charge-coupled device, MM: multi-mode fiber, SM: single-mode fiber.

Tumors were irradiated from the top at a treatment wavelength *λ*_*T*_ = 808 nm with a continuous wave diode laser (LuOcean Mini, Lumics, Germany) using 2 W/cm^2^ intensity, 1.2 cm illumination spot diameter and 5 min duration. GNR–PEG concentration and irradiation conditions were selected based on previous experiments [53]. After irradiation, tumors were allowed to cool down for another 5 min before continuing with the next steps.

Throughout the PPTT procedure, skin temperature was continuously monitored with an infrared camera (FLIR A35sc, FLIR Systems, USA). Tumor skin temperature was measured as the average in a region of interest (ROI) of circular area, centered around the pixel of maximum temperature with a fixed radius of 24 pixels (∼2 cm). The maximum ROI-averaged temperature for a mouse, reached at the end of the 5 min of irradiation, will be referred to as *T*_*ROI*_.

Mice were also monitored with a paw pulse-oximeter (MouseSTAT, Kent Scientific), recording heart rate and peripheral tissue oxygen saturation *SpO*_2_.

Diffuse optical data was acquired before and after PPTT, as described in the following sections and schematized in Figure 1A.

As a preventive measure, silver sulfadiazine (Silvederma spray, Laboratorios Aldo-Unión, Spain) was applied topically to all treated animals, once per day at 0, 24, 48 and 72 h after PPTT, to reduce skin burns and associated infections. After PPTT, mice were returned to their cages and tumor volumes were measured by palpation every 3 days.

Animals for short-term studies were sacrificed 24 h post-therapy, while animals for long-term studies formed two groups: (a) those sacrificed at a fixed time point of 16 *±* 1 days post-therapy for histological comparison and (b) those left for free survival, sacrificed when their welfare was compromised, to follow the evolution of tumor volume.

The long-term outcome of the therapy was analyzed through mouse survival time and progression-free survival (PFS) time, defined as the time to a 50% tumor volume growth after therapy.

In every case, at the end of the experiment, animals were sacrificed by cervical dislocation and the tumor and surrounding tissues were harvested and stored for posterior analysis.

### 2.6 Immunohistochemistry assays

After sacrifice, tumors and possible attached organs (kidney, spleen and/or skin) were extracted, weighed and cut in halves. In most cases, the skin was maintained attached to the tumors and used as a reference to orient the histological cuts. Tissues were fixed in neutral buffered formalin 10% for 24 – 48 h and processed for paraffin embedding. Specimens were dehydrated in series of ascending concentrations of ethanol, then cleared by serial xylene immersions, infiltrated in melted paraffin and embedded in hard paraffin wax.

Formalin-fixed paraffin-embedded (FFPE) mouse tissue samples were sliced at 3 – 5 µm and stained with hematoxylin–eosin (H&E), and with anti-cleaved caspase 3 (Asp175) rabbit polyclonal antibody (#9661, Cell Signaling Technology, USA) or the corresponding isotype control (ab172730) for the negative control.

Full images of immunostained sections were acquired with a slide scanner (NanoZoomer-2.0 HT C9600, Hamamatsu, Japan) at 20X magnification (resolution of 0.46 µm/pixel). The H&E staining was used to quantify cell death by necrosis in each section and the expression of cleaved caspase-3 (Casp3) indicated the cells that underwent apoptosis.

For the quantification, individual selection of each neoplastic growth was performed manually and each region of interest was analyzed using the pixel count, with 3,3’-diaminobenzidine (DAB) as staining for cleaved caspase 3 (positive pixels) and hematoxylin as counterstaining (negative pixels).

Area measurements were done manually using QuPath software [54]. White areas (tissue artifact/background and undefined spaces) were not included. All immunohistochemistry sample preparation, stainings and analysis were conducted by histopathologists from the Servei d’Histopatologia de l’Institut de Recerca Biomèdica de Barcelona (IRB, Barcelona).

### 2.7 Diffuse optical monitoring

A hybrid diffuse optical system (**Figure 1C**) combining DRS and DCS techniques was used for tissue optical and hemodynamic monitoring over the different steps of PPTT. This device has already been shown capable of retrieving absolute optical and hemodynamic parameters and GNR concentration in these pre-clinical models. Here we provide a brief description of the system and further details are available in the references [47, 48].

A rigid contact probe with a semicircular face of radius of ∼3 mm, containing the source and detector fibers for both techniques, was placed in contact with the hairless skin of nude mice over the tissue to interrogate and fixed in place with a manually adjustable holder.

#### 2.7.1 DRS system

The DRS system used a quartz tungsten-halogen broadband source (QTH model 66499, lamp model 6334NS, Oriel Instruments, Newport Corporation, USA), whose light was guided by a multi-mode fiber Y-bundle (400 µm core) to two locations on the face of the rigid probe. These two source fibers, together with six detector fibers (200 µm core), were utilized in two sets of source-detector pairs.

The first source, at the center of a semicircle of collectors, made equal source-detector separations (SDS) of 0.25 cm used for calibration measurements. The second source, located at a vertex of the semicircle, formed the second set of pairs, with SDSs ranging from 0.25 to 0.5 cm, used for probing different volumes of the tissue.

A spectrograph and integrated CCD camera (Acton Insight with 1340 *×* 400 pixel CCD PIXIS eXcelon 400B, Princeton Instruments, USA) obtained a wavelength-resolved measurement for each pair in the 550 – 1050 nm range.

#### 2.7.2 DCS system

The DCS system consisted in a long-coherence length 785 nm laser (DL785-120-S, CrystaLaser, USA) guided with a multi-mode fiber to the probe, four single-mode fibers (5.6 µm core) for light collection connected to single-photon counters (SPCM-ARQH-14, Excelitas Technologies, USA) and respective auto-correlator boards (Correlator.com, USA) to calculate intensity autocorrelation curves. An optical switch alternated between two sources fibers on the probe to measure twice as many SDS values in the same range as DRS (0.25 – 0.5 cm).

#### 2.7.3 Measurement protocol

The anesthetized mice were placed on a thermal bed were isolated from external ambient light in a cabinet designed for this purpose. The probe was placed over the tumor or shoulder of the mice, making sure the whole face was in contact with the skin. For each placement of the probe, fiber shutters alternated five times between DRS and DCS sources, synchronized with detector measurements. An extra DRS dark frame was acquired at the beginning of each repetition for background noise subtraction.

The probe was relocated five times over the tumor in different positions to account for spatial heterogeneity and three times over the shoulder on the same location.

For the study of GNR–PEG accumulation in tumors, measurements were performed right before injection and 24 h later. For the study of PPTT, measurements were performed immediately before and immediately after the therapy (including cooldown time). This is schematized in Figure 1A.

#### 2.7.4 Diffuse reflectance model for DRS analysis

Broadband DRS data was fitted using the delta-Eddington *P*_1_ approximation (*δ*-*P*_1_) to the radiative transport equation for a homogeneous medium and a pencil source, as detailed in reference [55]. Details for the model used can be found in Section 3.1 of the Supporting information.

Briefly, under this model the macroscopic behavior of light is be described by the absorption coefficient *µ*_*a*_, the scattering coefficient *µ*_*s*_ and two anisotropy factors for the scattering, that will be fixed to the standard values *g* = 0.9 and *γ* = 1.9 [56]. The usual relation 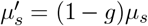 defines the probability of isotropic light scattering events, or reduced scattering coefficient. The diffused fluence rate Φ_d_(*ρ, z*) takes the form:

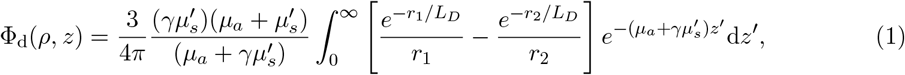

where *ρ* is the source-detector separation, *z* is the inward spatial dimension into the tissue, 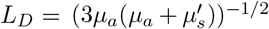 is the optical penetration depth and *r*_1_ and *r*_2_ are the distances to the effective isotropic real and image source points, respectively.

#### 2.7.5 DRS fitting algorithm

DRS data was fitted following the self-calibration method used in references [47, 48] and implementing an iterative fitting algorithm. Prior to fitting, the dark-subtracted signal of each collector, *I*_*j*_(*λ*) (*j* = 1, …, 6), was divided by the calibration measurement 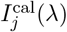 (same collection fiber, illumination through calibration source) in order to remove collector–skin coupling effects and the source spectrum factor. Later, this ratio was normalized to that of the previous collector in order to minimize source–skin coupling effects [57]. This can be summarized for collectors *j* = 2, …, 6 as:

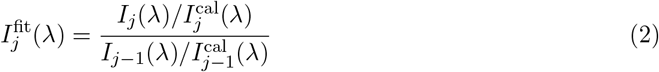

Thus, the quotient Φ_d_(*ρ*_*j*_, 0)*/*Φ_d_(*ρ*_*j*−1_, 0) was fitted to the five resulting normalized signals 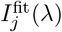 separately, assuming a homogeneous medium each time, to allow for tissue heterogeneity.

Effective Mie scattering was assumed for the spectral dependence of 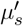, of the form 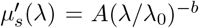, where the scattering amplitude *A* and the scattering power *b* were fitted experimentally and the reference wavelength *λ*_0_ = 785 nm was used [58, 59].

For the absorption coefficient *µ*_*a*_, linear contributions were considered from the main chromophores, Hb, HbO_2_, H_2_O and GNRs, using *µ*_*a*_(*λ*) = ∑_*i*_ *C*_*i*_*ϵ*_*i*_(*λ*), where *ϵ*_*i*_(*λ*) is the absorption coefficient per unit molar concentration for chromophore *i* and *C*_*i*_, its molar concentration. As an exception, water contribution is expressed as 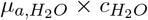 where 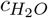 is the water fraction (dimensionless), and 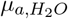, the absorption coefficient of pure water. The total hemoglobin concentration 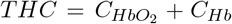 and tissue oxygen saturation 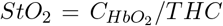 were fitted parameters together with *C*_*GNR*_ and 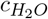. The full set of DRS fitted parameters was thus 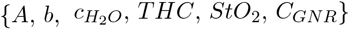.

An iterative fitting algorithm was tested and optimized with *in vivo* data through objective analysis of the residuals. In the chosen approach, each variable was fitted roughly in the wavelength range were they had a higher impact on the reflectance, while leaving the rest of parameters constant, using a non-linear least-squares algorithm in MATLAB software (Version 9.5.0.1067069 (R2018b) Update 4, The Mathworks, Inc.). In addition, the first derivative of the signal with respect to wavelength was exploited to improve the determination of 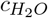 around the absorption peak at 970 nm, as this helped to decouple it from *C*_*GNR*_. The relative residuals of the fit were used to determine sufficient convergence or discard the data. The full description of the iterative method is reported in Section 3.2 of the Supporting information.

#### 2.7.6 Diffuse correlation model

For the DCS analysis, the diffusion approximation to the correlation transport equation was used to model the electric field autocorrelation function *G*_1_(*ρ, τ*), were *τ* is the time delay [30]. DCS measurements at separations *>* 2 mm have been shown to have acceptable error in the determination of *BFI* using diffusion theory and the so-called effective Brownian motion assumption, so it was not considered necessary to adapt this model for our short SDS values [60]. Moving scatterers determine the decorrelation rate of the electric field according to the Green’s solution to said equation:

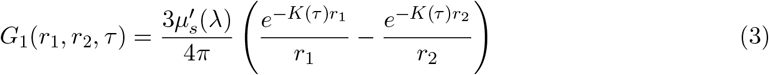

where 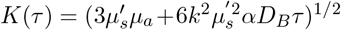, *k* is the light wavenumber in the medium, *α* is the fraction of scatterers that are not static and *D*_*B*_ is the scatterer diffusion coefficient (empirical Brownian model assumed). The blood flow index (*BFI*)is defined as the combination *αD*_*B*_ in *K*(*τ*).

Measured normalized intensity autocorrelation curves *g*_2_(*ρ, τ*) are related to normalized field autocorrelation curves *g*_1_(*ρ, τ*) through the Siegert relation: *g*_2_(*ρ, τ*) = 1 + *β* |*g*_1_(*ρ, τ*)|^2^, in which the coherence factor *β* accounts for the number of uncorrelated modes detected (0 *< β <* 1), expected to take the value 0.5 for the detection of single-mode non-polarized light [61].

#### 2.7.7 DCS fitting algorithm

The value of *β* was averaged from the first points of measured *g*_2_ curves to account for small deviations from the ideal value and *g*_1_ was then obtained through the Siegert relation, clipping *g*_2_ values *<* 1 due to noise. Any autocorrelation curve with *β <* 0.45 or *β >* 0.55 was discarded. The values used for *µ*_*a*_ and 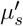 in Equation (3) were the fitted values for the corresponding SDS, so as to account for any possible depth heterogeneity. *BFI* was not fitted for the measurements where the DRS fits were discarded.

### 2.8 Statistical methods

#### 2.8.1 Growth model for tumor volume

To obtain the mean exponential growth rates for each experimental group, a linear mixed effects (LME) model was used to fit the log-transformed values of tumor volume over the time from irradiation. In particular, the fitted model was:

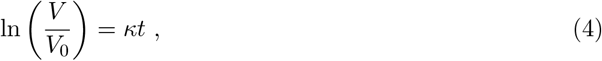

where *V* is the tumor volume, *V*_0_ = 1 cm^3^ is the pre-defined volume at the day of treatment, *t* the time after therapy and *κ* the growth constant. The model used considered null intercept (as the volumes were normalized to the initial value), a different growth constant for each group (fixed effect) and each mouse (random effect). In other words, the model also provided a growth model for each individual mouse. Pairwise tests between groups were conducted using Bonferroni-adjusted two-tailed Z-tests.

#### 2.8.2 Group survival and PFS analysis

Survival analysis was used for testing differences between groups in time-to-event outcome variables: mouse survival and PFS (50% growth threshold). Standard Kaplan–Meier analysis with pairwise log-rank test was used for pairwise comparisons between groups. Animals sacrificed at fixed time points were included in the analysis but right-censored from the day of sacrifice onwards. In the case of PFS, they were right-censored if they died before reaching the volume threshold [62].

#### 2.8.3 Relations between survival or PFS and continuous covariates

To test for the effect of different continuous variables on survival and PFS (in addition to group differences when appropriate), Cox proportional hazards models [63] were implemented for the hazard ratio *HR* between two mice, as ln(*HR*) = ∑_*i*_ *B*_*i*_∆*x*_*i*_, where *B*_*i*_ is the effect size of the difference in the covariate *i*, ∆*x*_*i*_, between the mice. The probability that an individual *j* suffers the event at a time *T*_*j*_ longer than another individual *k, T*_*k*_, is given by the so-called probabilistic index (PI): [64, 65]

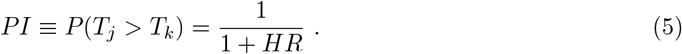

The fit residuals were checked for violation of proportionality of hazards by testing the correlation with time of Schoenfeld residuals [66].

#### 2.8.4 Software

Statistical analysis was conducted using R Statistical Software (v4.0.0, R Core Team 2020), using base packages and “lme4” (v1.1.23), “performance” (v0.9.1), “multcomp” (v1.4.13), “survminer” (v0.4.7) and “survival” (v3.2.3) packages; and MATLAB using base packages and the “Linear Deming Regression” package (v1.2).

Other details about statistical tests are reported in Section 4 of the Supporting information.

### 2.9 Light transport and heat diffusion simulations

Monte Carlo (MC) light transport and heat diffusion (HD) simulations were carried out with MCmatlab open software [67] in MATLAB. For both, the same 101 *×* 101 *×* 101 square voxel mesh and media geometry were employed to represent a cube of 5 *×* 5 *×* 5 cm dimensions. A homogeneous medium was used to represent tissue, using typical thermal properties [24, 68] and the optical coefficient values measured experimentally to simulated each treated tumor. A layer of single voxels on the illumination face was optically defined as air (null absorption and scattering) to simulate reflections back into the tissue. For the investigation of therapy dynamics and the extrapolation of GNR accumulation conditions beyond the measured values, tumor *µ*_*a*_ and 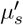 values were varied. The complete list of parameter values used for simulation can be found in Section 5 of the Supporting information.

Light transport simulations used 10^7^ photon packets incident in a 1.2 cm diameter beam (imitating experiments) with a top-hat profile, shone perpendicular to the top surface, with escaping boundary conditions in all faces.

A uniform initial temperature of 35°C and constant-temperature boundary conditions were assumed for heat diffusion simulations. The only heat source considered was the absorption of the therapy light, based on the MC derived fluence rate at each position in the tissue. The cooling of the tissue surface through the contact with outside air was neglected. The irradiation time and power were varied for the extrapolation of therapy conditions.

The mean temperature in the region of interest by the end of the 5 min of irradiation, 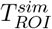, was calculated in analogy to experiments using the output of the simulation. The temperature of the voxels in the first layer of tissue was averaged inside a circle of 9 mm radius, centered on the illumination point.

## 3 Results

### 3.1 GNR–PEG biodistribution

The biodistribution of GNR–PEG 24 hours after injection (21 nM, 100 µl, 23 – 26 g mice weight), assessed by ICP-MS analysis of blood and tumor (*n* = 14), as well as kidney, spleen and liver (*n* = 11) is shown in **Figure 2** in terms of (a) the fraction of the injected dose (%ID) accumulated and (b) the average GNR concentration per gram of tissue (mol/g).

**Figure 2:**
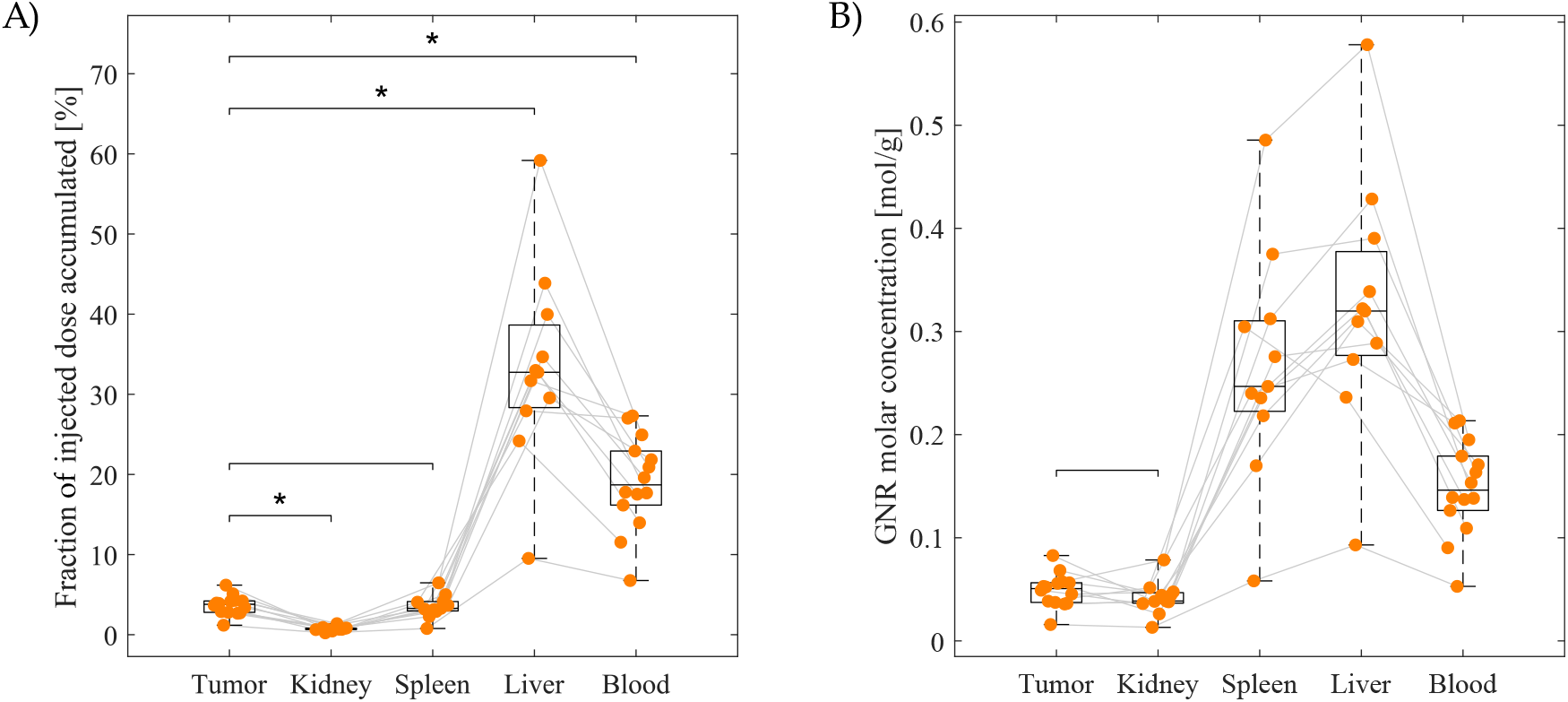
Quantification of GNR–PEG biodistribution. (A) Accumulated dose fraction and (B) concentration of GNR–PEG, in tumor, blood (*n*=14), kidney, spleen and liver (*n*=11) 24 h after injection. The boxes indicate the medians and interquartile ranges, the whiskers show the full ranges, the brackets indicate the pairwise tests conducted (Wilcoxon signed-rank test with Bonferroni correction, *: *p<*0.05) and the grey lines match data points from the same mouse.

The largest collector of GNR–PEG was the liver, with 33 *±* 3%ID (median *±* standard error of the mean, SEM)). In second place, 19 *±* 2%ID was still circulating in the blood by 24 h. Gold content in the tumor was found to be 3.8 *±* 0.3%ID, lower than in the liver (Bonferroni-corrected *p*-value *p <* 10^−2^) and the blood (*p <* 10^−3^) and greater than in the kidney (0.68 *±* 0.08 %ID, *p <* 10^−2^).

The GNR concentration in the tumor, of (5.1 *±* 0.4) *·* 10^−2^ mol/g tissue, was not significantly different than in the kidney where the tumor was grafted, of (3.9 *±* 4) *·* 10^−2^ mol/g tissue.

### 3.2 GNR–PEG toxicity profile

During the six-month period of evaluation of toxicity, animals did not show signs of discomfort or changes in behavior according to welfare guidelines.

The histological examination of the main organs six months after the administration of the usual working dose of GNR–PEG found hyperplasia in Langerhans islets, hepatic glucogenosis and different degrees of hepatic lipidosis.

Biochemistry and hematology analysis results showed no statistically significant differences between animals injected with GNR–PEG and PBS. Complete blood test results are reported in Section 2 of the Supporting information.

### 3.3 Tumor volume evolution and mouse survival after PPTT

PPTT outcome with fixed therapy conditions was evaluated on thirty mice in the GNR–PEG group (*n*=30) and twenty-seven mice in the PBS group (*n*=27), while nineteen mice receiving no injection or irradiation were used as the control group (*n*=19). All groups and numbers of animals in PPTT experiments are detailed in Table 1. For the evaluation in the long term, a cohort of thirty-five mice (*n*=35) among the three groups were followed up after therapy and sacrificed when their welfare was compromised (indicated in the table as “free survival”), while twenty (*n*=20) were sacrificed at set time points after therapy for *ex vivo* assays (indicated as “fixed survival”).

**Table 1:** Number of mice in each experimental cohort and group.

| Survival conditions |  | Number of mice |  |  |  |
| --- | --- | --- | --- | --- | --- |
|  |  | GNR-PEG | PBS | Control | Total |
| Long-term | Free survival: 9–27 days | 14 | 11 | 10 | **35** |
|  | Fixed survival: 15–17 days | 7 | 8 | 5 | **20** |
| Short-term | Fixed survival: 1 day | 9 | 8 | 4 | **21** |

During laser irradiation, the ROI-averaged skin temperatures *T*_*ROI,t*_ raised with varying slope, as shown in **Figure 3A**. The average *T*_*ROI*_ values (after 5 min of irradiation) were significantly different between the irradiated groups (*p <* 10^−3^). The values reached were 46.4 *±* 0.6 °C (mean *±* SEM) for the PBS group (11.4 *±* 0.6 ° increase) and 58.0 *±* 0.8 °C for the GNR–PEG group (23 *±* 0.8 °C increase).

**Figure 3:**
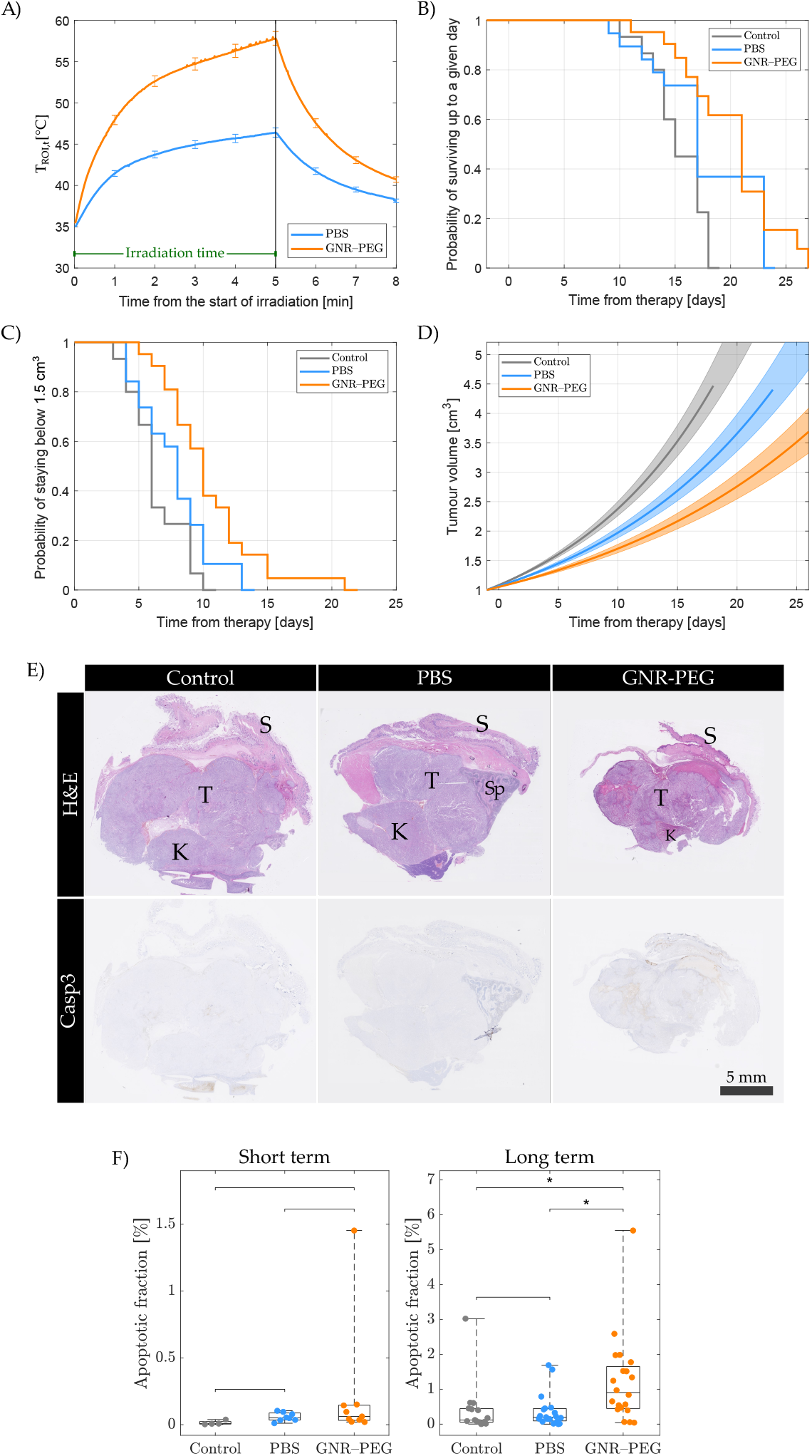
Outcome assessment with standard methods. (A) Mean (*±* SEM) skin temperature averaged over the ROI (*T*_*ROI,t*_), during 5 min of irradiation and 3 min of cooling (*N*_PBS_=27, *N*_GNR_=30). (B) Survival Kaplan–Meier plot (long-term cohorts: *N*_control_=15, *N*_PBS_=19, *N*_GNR_=21). (C) PFS Kaplan–Meier plot (long-term cohorts). (D) Fitted LME model tumor volume curves (long-term cohorts). Lines show group estimates (up to maximum survival day) and shaded regions indicate SE range in fit parameters. (E) Representative images of H&E and Casp3 staining (20X magnification) 15 – 17 days after therapy. T: tumor, K: kidney, S: skin, Sp: spleen. (F) Casp3 staining results at 1 day (short term, *N*_control_=4, *N*_PBS_=8, *N*_GNR_=9) and 15 – 17 days after PPTT (long term, *N*_control_=5, *N*_PBS_=8, *N*_GNR_=7). Boxes indicate medians and interquartile ranges, whiskers show full ranges and brackets indicate pairwise tests conducted (Wilcoxon rank-sum test with Bonferroni correction, *: *p<*0.05).

A Kaplan–Meier plot for mouse survival is shown on **Figure 3B**. Survival half-lives were of 15 days for controls, 17 days for the PBS group and 21 days for the GNR–PEG group. Survival was statistically significantly different between the GNR–PEG group and control mice (*p* = 0.03), indicating extension of survival by PPTT.

The Kaplan–Meier plot for PFS is shown on **Figure 3C**. Two animals in the GNR–PEG group were sacrificed before reaching the 50% growth threshold (right-censored at their time of death). PFS half-lives were of 6 days for controls, 8 days for the PBS group and 10 days for the GNR–PEG group. The probabilities of staying under the threshold were significantly different between all groups (*p <* 10^3^). This meant that the tumor growth was delayed by laser irradiation and further delayed with the contribution of GNRs.

Group estimates of volume progression (LME model fitted to volume data) are presented in **Figure 3D**. Multiple comparison showed that the growth for each group was statistically significantly different to the rest (Control – GNR–PEG *p <*10^−3^, Control – PBS *p*=0.017, GNR–PEG – PBS *p*=0.049). The growth constants for each group (estimate *±* SE) were found to be 0.079 *±*0.005 day^−1^ for controls, 0.062 *±* 0.004 day^−1^ for the PBS group and 0.048 *±* 0.004 day^−1^ for the GNR–PEG group (see Equation (4)). The individual tumor growth models are shown in Section 6 of the Supporting information. The residuals of the fit were normally distributed (Shapiro-Wilk test, p*>*0.05).

### 3.4 Triggered cell death pathways in treated tumors

Representative tumor slices stained in H&E and Casp3 assays for the estimation of necrotic and apoptotic tumor fractions are shown on **Figure 3E**. Assays in mice sacrificed 1 day after PPTT (*n* = 21, **Figure 3F**, left) and 15 – 17 days after PPTT (*n* = 20, Figure 3F, right) found that for the GNR–PEG group the apoptotic fraction was higher in the long term than in the short term (*p* = 0.03). In addition, in the long term, there was a higher apoptotic fraction in the GNR–PEG group with respect to the PBS group (*p* = 0.04). There were no significant differences between groups of mice sacrificed in the short term.

The results of the measured necrotic fraction are reported in Section 7 of the Supporting information, with no significant differences between groups in either the short or long term.

### 3.5 Relationship between hemodynamic tumor properties and GNR accumulation

Diffuse optical measurements in tumors before and 24 h after the injection of GNR–PEG (*n* = 40) were analyzed to relate the accumulation of GNRs in a tumor to its hemodynamic properties (timeline schematized in **Figure 4A**). Tests revealed a statistically significant exponential dependence of the delivered GNR dose with pre-injection 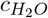 (LME, *p <* 10^−3^). The model (**Figure 4B**) found an intercept for the logarithm of GNR concentration of 3.4 *±* 0.1 nM and slope of 0.5 *±* 0.1 nM (estimates *±* SE).

**Figure 4:**
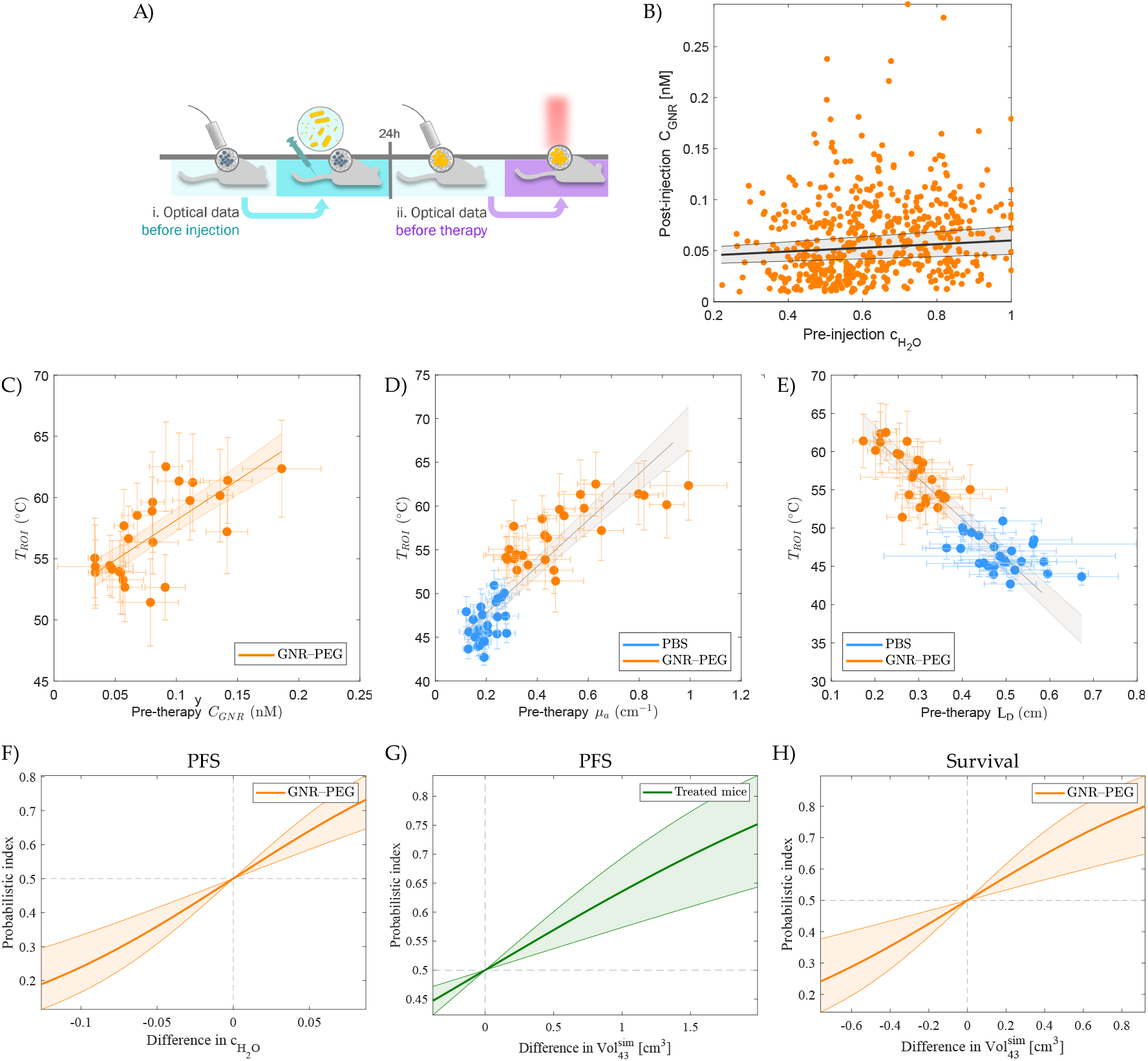
Associations between relevant therapy markers and diffuse optical data obtained before NP injection or before therapy. (A) Measurement timeline. (B) Fitted LME model for the relation between *C*_*GNR*_ 24 h post-injection and pre-injection 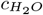 (see 2.8 for details of the model). Each point is a measurement on a tumor (five per mouse with sensitivity cutoff, *n*=40). The shaded region indicates the SE range in fit parameters. (C-E) Relationships between *T*_*ROI*_ values (IR thermography) and pre-therapy tumor optical properties (DRS), overlayed with Deming regression models: *C*_*GNR*_ (only GNR–PEG group), *µ*_*a*_(*λ*_*T*_) and *L*_*D*_ (*λ*_*T*_) (*n*_GNR_=30, *n*_PBS_=27)). Each point is the median for a mouse with SD bars. The shaded regions indicate the SE range. (F-H) Probabilistic indices for Cox models relating pre-therapy optical measurements/simulations to outcome variables for the observed range of absolute differences between subjects: PFS dependence on 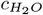 for the GNR–PEG group, PFS dependence on 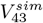 for all treated mouse and survival dependence on 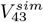 for the GNR–PEG group (long-term cohorts: *N*_PBS_=19, *N*_GNR_=21). The shaded regions indicate *e*^−1^ confidence intervals.

### 3.6 Prediction of the temperature reached during therapy

For the GNR–PEG group, *T*_*ROI*_ values were correlated with the *C*_*GNR*_ (*p <* 10^−3^) measured right before irradiation. A Deming regression (**Figure 4C**) found an intercept of 51.6 *±* 0.9°C and slope 65 *±* 10°C/nM. In addition, *T*_*ROI*_ was also correlated with *THC* within each treated group (*p <* 10^−3^, figure S3 in Supporting information, Section 8). Deming regression revealed for the GNR–PEG group an intercept of 50 *±* 2°C and a slope of 0.05 *±* 0.02°C/µM and, for the PBS group, an intercept of 40 *±* 2°C and a slope of 0.10 *±* 0.03°C/µM.

Generalizing the approach to all treated mice (PBS and GNR–PEG groups), the hypothesis of a positive relationship between *T*_*ROI*_ and pre-irradiation *µ*_*a*_(*λ*_*T*_) was corroborated (*p <* 10^−3^). The Deming regression (**Figure 4D**) revealed an intercept of 40 *±* 1°C and a slope of 30 *±* 3°C cm.

*T*_*ROI*_ was compared with the pre-therapy *L*_*D*_(*λ*_*T*_) to evaluate the effects of the characteristic light diffusion length. The Deming regression (**Figure 4E**) revealed an intercept of 70 *±* 1°C and a slope of 50 *±* 4°C/cm (*p <* 10^−3^).

### 3.7 Pre-therapy prognostic models

The relations between therapy outcome variables (survival and PFS) and all optical parameters measured before therapy were evaluated in the groups of treated mice. Cox models showed that only a higher 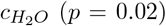 was associated to a retarded volume progression (longer PFS) for mice in the GNR–PEG group. The fitted slope for the log(HR) (*±* standard error, SE) was −12 *±* 5. The PI for this model (Equation 5) is shown in **Figure 4F**. For instance, a GNR–PEG mouse with a 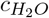 value higher by 0.1 than another mouse had a 76.1% [66.7%, 83.5%] estimated probability [*e*^−1^ confidence interval]) of having a longer PFS. No other raw optical or hemodynamic parameter was significant in Cox models.

Mouse survival was not associated with optical parameters but it was affected by the mouse body weight at the day of therapy (24.7 – 41.3 g, *p* = 0.006). The fitted estimates for the log(HR) were −0.6 *±* 0.5 for the PBS group, −1.3 *±* 0.5 for the GNR–PEG group, both with respect to controls, and −0.18 *±* 0.07/g for the slope with weight. The probabilistic index for the fitted Cox model can be found in Figure S4 in Supporting information, Section 9.

### 3.8 Pre-therapy prognosis based on simulations

Simulated temperature maps were validated against experimental data, finding 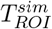 was linearly correlated with *T*_*ROI*_ with an intercept of -0.4 *±* 4 °C and slope of 1.02 *±* 0.09 (*p <* 10^−3^, data not shown). Then, the volumes of voxels that reached the significant temperature thresholds of (a) 43°C (onset of irreversible damage), 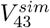, and (b) 55°C (coagulation in a matter of ∼ 10 s), 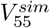, were used as surrogate simulated parameters for the estimation of therapy outcome.

Both 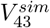 and 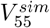 were significantly associated with the PFS of all treated animals in Cox models (respectively *p* = 0.03 and *p* = 0.048). In the case of the latter, only the mice with 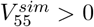 were considered, leaving out nine PBS and one GNR–PEG mice.

The estimated slope for the log(HR) *vs*. 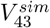 was −0.6 *±* 0.3/cm^3^. **Figure 4G** shows the PFS PI as a function of 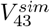. A treated mouse had a 63.8% [57.3%, 69.3%] probability of having a longer PFS than a mouse with a 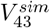 value lower by 0.5 cm^3^. On the other hand, the estimated slope for the log(HR) *vs*. 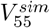 was *™*1.4 *±* 0.8/cm^3^. A treated mouse had a 67.2% [58.3%, 75.0%] probability of having a longer PFS than a mouse with a 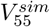 value lower by 0.5 cm^3^.

Furthermore, the survival of mice in the GNR–PEG group was significantly related to 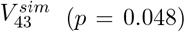, with an estimated slope of *™*1.5 *±* 0.8/cm^3^ for the log(HR). **Figure 4H** shows the survival PI as a function of 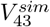. A mouse in the GNR–PEG group had a 67.9% [58.2%, 76.4%] probability of having a longer survival time than a mouse with a 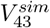 value lower by 0.5 cm^3^.

### 3.9 Extrapolation to other therapy conditions using simulations

The figures of merit chosen to estimate the outcome through the output temperature field were 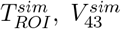 and 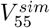. **Figure 5** summarizes, for a general comparison between groups, the extrapolation of the figures of merit to different therapy conditions. We have used typical tumor absorption coefficient values for each group (*µ*_*a*_= 0.2 cm^−1^ for PBS group, *µ*_*a*_= 0.4 cm^−1^ for GNR–PEG group) and identical scattering properties for both (*µ*_*s*_= 80 cm^−1^ and *g* = 0.9).

**Figure 5:**
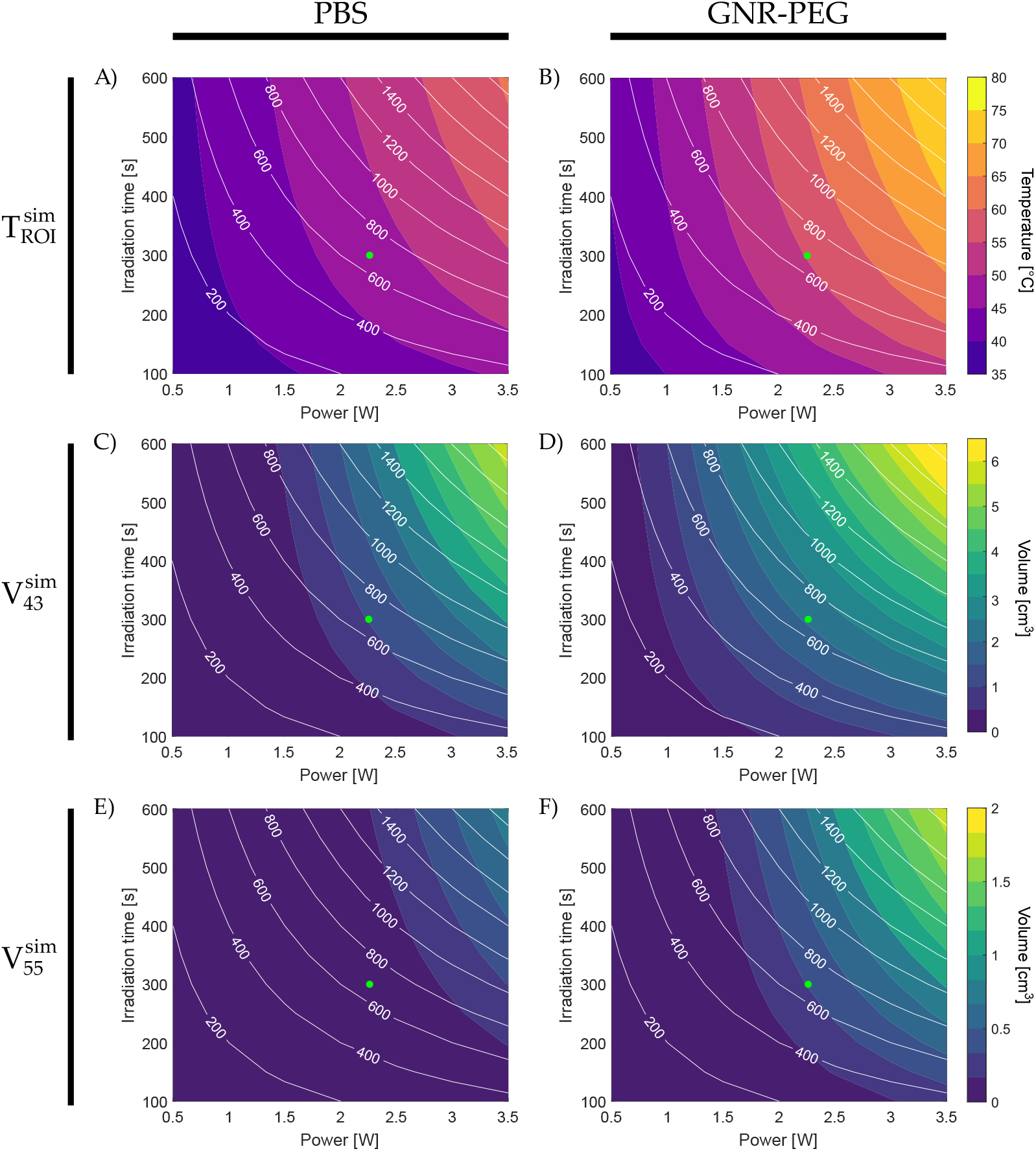
Extrapolation of simulated parameters 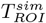 (top), 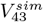 (middle) and 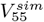 (bottom) to different values of irradiation power and time for typical tumor optical properties with PBS injection (left column) and with 21 nM GNR–PEG injection (right column). Level curves for the total energy delivered are shown in white in units of joules. Green points mark the therapy conditions used in our experiments (2.26 W power and 300 s irradiation time).

### 3.10 Hemodynamic changes after therapy

Immediately after therapy, the skin over the tumors was visibly white, especially for GNR– PEG mice, with no open wounds due to burns. Post-therapy tumor optical data was successfully acquired and fitted using the same method as for pre-therapy measurements. The variations in hemodynamic parameters for each group are reported in Section 10 of the Supporting information, with decreases in 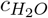, *THC, StO*_2_ and *BFI*.

The values of tumor 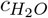 and *BFI* measured after therapy (timeline schematized in **Figure 6A**) were related to *T*_*ROI*_ values for the treated mice, as shown in **Figure 6B-C**. 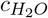 and *T*_*ROI*_ were correlated (*p <* 10^−3^) with an intercept of 1.3 *±* 0.1 and a slope of 0.013*±*0.002/°C, while the log(*BFI/*10^−8^ cm^2^/s) was correlated (*p <* 10^−3^) with an intercept of 10.1 *±* 0.9 and a slope of −0.20 *±* 0.02/°C.

**Figure 6:**
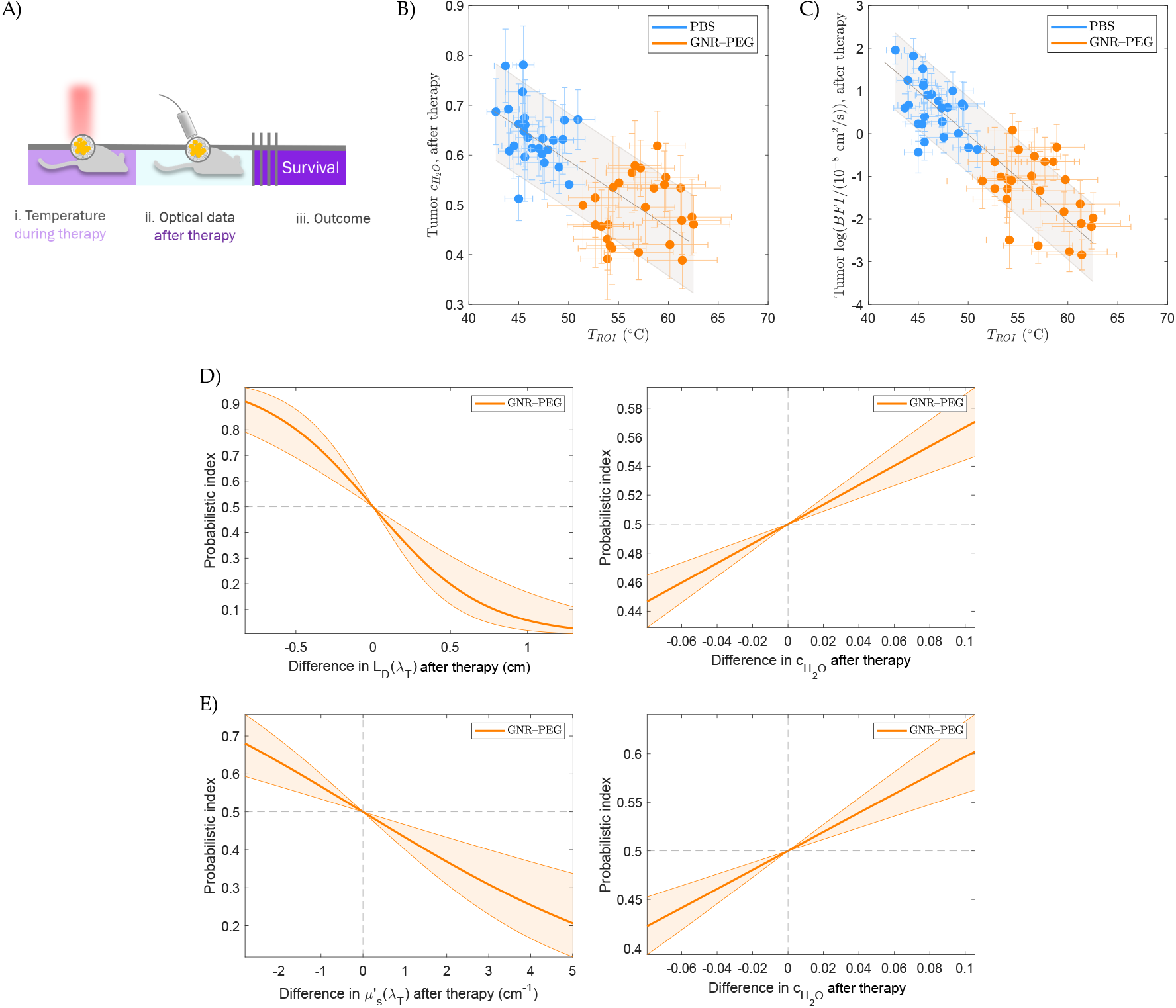
Associations between diffuse optical data obtained after therapy, *T*_*ROI*_ and outcome variables. (A) Measurement timeline. (B-C) Post-therapy tumor 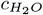 and log(*BFI*) versus *T*_*ROI*_ values with fitted Deming regression models (*n*_GNR_=30, *n*_PBS_=27)). (D) Dependence of the probabilistic index for mouse survival (bivariate Cox model) on post-therapy *L*_*D*_ (*λ*_*T*_) and 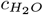 for the GNR–PEG group (long-term cohorts: *N*_GNR_=21), plotted over the range of observed pairwise differences between subjects. Shaded regions indicate *e*^−1^ confidence intervals. (E) Dependence of the probabilistic index of PFS (bivariate Cox model) on post-therapy 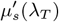 and 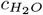.

### 3.11 Post-therapy prognostic models

The use of skin temperature information allowed to find a signification effect of *T*_*ROI*_ on volume progression (*p* = 0.02) in a Cox model for the PFS of treated animals. The fitted slope of the log(HR) *vs. T*_*ROI*_ (*±* standard error) was −0.07 *±* 0.03/°C.

Tumor optical and hemodynamic parameters measured after therapy were then used as additional information to predict the outcome of treatment in Cox models for mouse survival and PFS.

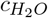 and *L*_*D*_(*λ*_*T*_) were significant parameters in a compound model for the survival of mice in the GNR–PEG group (*p* = 10^−3^ and *p* = 0.009, adding each variable in steps). The estimated slopes for the survival log(HR) were −23 *±* 9 for 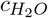 and 1.4 *±* 0.6/cm for *L*_*D*_(*λ*_*T*_). Moreover, 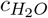 and 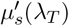 were significant in a compound Cox model for the PFS of mice in the GNR–PEG group (*p* = 3 *·* 10^−3^ and *p* = 0.04). The estimated slopes for the PFS log(HR) were −16 *±* 5 for 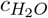 and 0.5 *±* 0.3 cm for 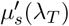. Figure **6D-E** shows the dependence of the survival and PFS PI for this group on optical parameters.

## 4. Discussion

This work consisted in the development and demonstration of a comprehensive set of techniques for the identification of optical and hemodynamic markers that enable the personalization of PPTT. It aims to address one of the main obstacles in the clinical translation of the therapy, which is the heterogeneity in the response to treatment, a challenge common to all cancer therapies and central in the clinical research of the management of the disease. Our toolbox gathers diffuse optical techniques and light and heat transport simulations to work in synergy with standard methods in nanomedicine and oncology in order to relate tumor tissue properties with NP delivery, hyperthermia, treated volumes and therapy outcome. These findings could then be used as rules to adjust therapy parameters based on individual tumor properties to optimize the response to the therapy and reduce the variability observed.

In the first place, therapy safety and efficacy were evaluated to establish its applicability and allow the assessment of our personalization toolbox. Moreover, evidence of good NP biodistribution and long-term toxicity profiles are considered prerequisites for the application and translation of any particular NP formulation in a PPTT protocol, as NP shape, surface chemistry and hydrodynamic size can govern their elimination routes, circulation and retention times, and thus determine their safety and efficiency as mediators of PPTT [69–72].

For our particular formulation, the GNR–PEG concentration delivered to the tumor by 24 h after injection was in line with literature reviews [73] and similar to the concentration in the kidney, in agreement with two-photon microscopy measurements previously published for this model [50]. While the GNR–PEG were primarily captured by the liver and spleen, blood content at 24 h implies a long circulation time, which favors accumulation in the tumor [71]. These results are typical for NPs larger than 6 – 8 nm, as they cannot be filtered by the renal glomerulus, and thus are captured by the mononuclear phagocytic system (mainly the liver, spleen and lymph nodes) or extravasate into tumors [71, 74]. Fecal elimination route via hepatobiliary excretion has been described for the same GNR–PEG [75], confirming this effect.

In the long-term evaluation of toxicity, there were no relevant signs of chronic nanomaterial toxicity in the studied organs, with all histopathological findings linked to age and the obese phenotype of animals fed in ad libitum diets. Likewise, a different study with a five times higher dose of same GNR–PEG obtained similar results in C57Bl6J immunocompetent mice [75], supporting the biosafety profile of these nanoparticles.

Therapy efficacy in the GNR–PEG group was characterized by the extension of mouse survival with respect to control animals (figure 3B) and the retardation of tumor growth with respect to the PBS group and controls (figure 3C). The treatment led consistently to a decrease in tumor growth rates and not a reduction in tumor volume, showing that some areas of the tumors were untreated and kept growing, which was confirmed by histological examination. Similar effects have been reported in small animals in PPTT treatment of tumors of similar size [76, 77], in contrast with smaller tumors which are easily eliminated (for example, [78–84]). The large dispersion in PFS times within the treated groups supports the need of individual data for the personalization of the treatment protocol.

Cell death pathways were also explored in our experimental groups to gather additional information of the underlying biological mechanisms activated by PPTT. The signaling pathways of both apoptosis and necrosis can be triggered by mild hyperthermia [20, 85, 86], and their extent will vary depending on PPTT conditions. Given the nature of both pathways (regulated *vs*. uncontrolled response), activating apoptosis mechanisms for tumor cell death has always been preferred [20], as necrosis can paradoxically result in increased tumor cell proliferation or migration or an excessive immune response [87]. Even though no differences were observed in neither necrosis nor apoptotic markers at early stages in our experimental groups, the significant increase in caspase-3 observed in the GNR-PEG group two weeks after treatment, together with an unchanged necrosis profile, suggests a PPTT-induced activation of the apoptotic cell death pathway, as reported by others [88–91]. This sustained expression of apoptosis effectors might explain the significant delay in tumor growth observed in GNR-PEG mice; however, other mechanisms, such as the immunogenic cell death [92, 93], have also been suggested to be triggered by PPTT and could also contribute to the reduction of the tumor.

Once the safety, efficacy and heterogeneity in the response to therapy were established, we studied the capabilities of NIRS techniques to acquire personalized tumor information relevant to the progress and success of the therapy. Optical measurements taken pre-injection, pre- and post-irradiation allowed to correlate optical and hemodynamic properties with (i) NP accumulation in the tumor, (ii) the temperature rise during irradiation and (iii) the outcome of the therapy.

NP accumulation in tumors has been extensively attributed to the enhanced permeability and retention (EPR) effect [8], observed in rodents, humans and other mammals. This is the result of the fenestration of tumor neovasculature, chaotic and reduced blood flow, deficient vascular smooth muscle, malfunctioning endothelial cells and lack of functional lymphatic drainage [9, 43]. However, these characteristics are highly variable among tumors, modulating the EPR effect [44, 94]. It has even been proposed that its degree on each individual tumor should be quantified before deciding on a treatment [95].

Here, we hypothesized that the variations in the concentration of GNR–PEG measured in our tumor models 24 hours after injection responded to blood content and/or blood flow values at the moment of injection. Such a relation between hemodynamics and NP accumulation could then be used as a rule to adjust the injected dose for a desired result.

Our model showed that tumor *c*_*GNR*_ was only associated with 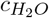, a parameter that could be indicating the volume of blood, lymph and other fluids implicated in the transport of GNRs. The mechanisms underlying this relation could be subject of further research of the microvascular characteristics of each tumor. Although there was a high variability observed among mice beyond this effect, which could prevent a precise control of the accumulated dose in this particular case, our methods show the potential to adjust the delivery and can still compensate for it in the following steps of the personalization of therapy.

The determinant factor for the therapeutic effect of PPTT is the evolution of tissue temperature in the volume of the tumor. Thus, its precise non-invasive monitoring during irradiation represents a powerful tool to evaluate the efficacy of treatment in real time. This has been demonstrated during photothermal therapy with a variety of MRI techniques [46, 78, 96–98] and thermometry through photoacoustic imaging (PAI) (*ex vivo* [99, 100] and in a subcutaneous tumor [101]). However, both technologies are limited by artifact formation and non-uniform temperature resolution [99, 102].

Furthermore, MRI costs are elevated and PAI sensitivity is dependent on tissue optical properties. For these reasons, IR skin thermography still represents a convenient monitoring tool for PPTT, and we have shown here that it can convey relevant information. We observed that a higher *T*_*ROI*_ was associated with a longer PFS in treated mice, and can thus be considered a prognostic marker obtained by the end of therapy.

However, temperature measurements can only allow one to modulate or stop the therapy in real time (when heat has already been applied), whereas a personalized irradiation protocol can benefit from a prediction of the temperature rise, allowing to choose irradiation parameters like the time or power before application. This is one of the advantages of NIRS monitoring, as demonstrated in this work.

Light-to-heat conversion in a lossy medium is linearly related to the medium’s light absorption probability, so a positive correlation was expected between tumor temperature, pre-treatment *µ*_*a*_(*λ*_*T*_) and the concentrations of chromophores in the tissue, particularly of GNRs. Indeed, this was evidenced with the linear correlations between *T*_*ROI*_ and *C*_*GNR*_ (GNR–PEG group, figure 4B), the *THC* of each treated group and *µ*_*a*_ (figure 4C). These optical parameters are thus simple pre-treatment indicators of the skin temperature rise after 5 min of irradiation.

The third aspect to be covered by our toolbox was the prognosis of therapy. In a clinical application, optical measurements could be used as additional information to tailor the treatment or stratify the response and choose to re-treat a tumor or perform a different procedure. This resembles the methodology applied to other therapies like PDT [36, 38, 41] and chemotherapy [32].

In our model, the only pre-therapy indicator of a slower tumor growth for mice in the GNR–PEG group was 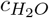 (figure 4F). Furthermore, this parameter was also the best post-therapy marker for both survival and PFS, indicating a high relevance of tumor water content for response, even when hyperthermia is known to cause dehydration [103, 104]) and post-therapy 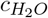 was inversely correlated with *T*_*ROI*_ in our experiments (figure 6A). Moreover, post-therapy models were better predictors of response, incorporating other variables like *L*_*D*_(*λ*_*T*_) and 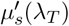. The changes in these parameters after hyperthermia were also in agreement with literature reports and related to temperature reached. An increase in 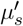 is caused by protein denaturation and loss of organization [105–108], which means a reduction in *L*_*D*_.

The next layer of modeling consisted in the use of the measured optical information in physical simulations, which expanded the functionality and capabilities of our toolbox. They allowed us to understand the physical mechanisms behind the skin temperature values observed and the treatment efficacy. Although MC and HD simulations have been used in the past to estimate the effects of PPTT in tissue (for example, [98, 109–115]), standard values for the optical properties have always been used, not specific to each tumor. In this work, we conducted an experimental validation of simulations on a mouse-to-mouse basis by imitating the processing of IR images. We achieved a very good correspondence between *T*_*ROI*_ and 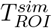 despite using fixed values for the index of refraction and thermal properties, which vary with water concentration [59, 116, 117]. We then identified the dominant effects on the evolution of temperature and found improved prognostic parameters of tumor growth. These findings enabled the extrapolation to other therapy conditions, which is an essential tool for the design of personalization protocols.

The search for new markers of treatment efficacy was based on exploiting personalized physical modeling. The simplest model for an estimation of the treated volumes is to consider the volume of tissue that reaches certain temperature thresholds during therapy (irradiation and cooling times) in the temperature map output of simulations. Although the time spent at an each temperature is important for the amount of damage produced in a portion of the tissue and it is considered in most damage models (such as CEM43 or Arrhenius models) [118], a simplistic approach like this one proved to be sufficient for useful results and avoids depending on a large number of physical properties that cannot be easily measured and vary largely across tissue types.

We chose the values of 43°C and 55°C as thresholds for our damage approximation based on the physiological effects of temperature on tissue. Whereas hyperthermia above 37°C and below 43°C induces reversible changes in biological tissues (increase in diffusion rates across membranes, synthesis of heat-shock proteins, blood flow, oxygenation and a reduction in pH) [16, 118–120], above 43°C, the irreversible unfolding and aggregation of protein structures start to occur, at rates depending on the tissue composition and the excess temperature. The persistence of elevated temperatures leads to an accumulation of damage and tissue deterioration. By 55°C, cell death is produced in a matter of minutes through the denaturation of proteins and DNA, the arrest of blood flow and thus the rapid decrease in oxygenation, and by 60°C, tissue coagulation is induced instantaneously. In this way, 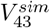 was used as an estimation of the volume of tissue with some level of thermal damage, while 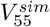 was an approximation for fully treated tumor portions, nonviable after hyperthermia.

Both 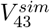 and 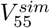 (simulated for each tumor) were significantly associated with the PFS of all treated animals (PBS and GNR–PEG groups together, figure 4G). This finding served both as a validation of our approximation of the treated volume and as a confirmation of improved pre-therapy prognostic markers of tumor growth. We note that they describe the outcome for all mice, and were not limited to the GNR–PEG groups, as opposed to what we had obtained using only optical measurements. Our results also show that in the PPTT temperature ranges it is an acceptable approximation to neglect air convection, perfusion cooling or the dependence of tissue properties on temperature.

Moreover, we investigated what were the effects of the heat generation profiles within the tissue on the skin temperature and treatment volumes. Experimentally, we found a negative correlation between *T*_*ROI*_ and the optical penetration depth *L*_*D*_(*λ*_*T*_) (figure 4D), which is a measure of how deep photons travel into the tissue (see Equation (1)). In a first interpretation, this suggests that a shallower penetration of light will cause heating closer to the skin. Later, independent sampling of *µ*_*a*_ and 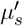 values in simulations (over experimental ranges) showed *µ*_*a*_ had a predominant effect on 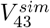 and 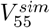, and although a higher *µ*_*a*_ led to a lower *L*_*D*_, the treated volumes increased due to the larger total amount of heat generated. This means the heat diffused deeper and achieved treatment temperatures in larger volumes of tissue despite the low penetration. See Section 11 of the Supporting Information for more details.

Finally, we extrapolated the treatment conditions to other powers, irradiation times and *µ*_*a*_(*λ*_*T*_) values. We demonstrated with the simulations the effect of delivering the same total energy at different rates (white level curves in figure 5). This had substantial implications for 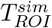 (figures 5A and B) and 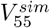 (figures 5E and F), where higher powers meant the energy concentrates momentarily where light fluence is highest, reaching higher temperatures before it diffuses. On the other hand, for 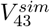 (figures 5C and 5D), following any energy level curve, the effect of power was important for values lower than 2 W, whereas there was little increase in affected volume for higher powers. This indicates that shorter irradiation times with higher powers are useful for treating larger volumes 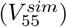 without affecting much larger regions of healthy tissue 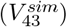.

Furthermore, the comparison between typical PBS and GNR–PEG *µ*_*a*_(*λ*_*T*_) values (left and right columns) showed that mice with no GNRs needed a much larger delivered energy to reach similar treated volumes as the GNR–PEG group. For example, a mouse in the PBS group would need to receive 1.5 – 2.5 times the energy to reach 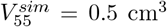. This would mean in turn 1.1 – 1.6 times the 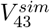 value than for the GNR–PEG case (depending on the conditions used, within the ranges shown in the figure). This illustrates the role of GNR in the confinement of thermal effects and sparing healthy tissue from being affected. Moreover, this effect was further seen over a larger GNR accumulation range in the extrapolation of tumor *µ*_*a*_(*λ*_*T*_) values (see Section 12 of the Supporting information).

Our methods and simulations have thus been shown to provide relevant markers for the personalization of PPTT and insight into the variability in response observed among tumors. What enabled this was primarily the possibility to measure individual tumor characteristics while simultaneously measuring the delivered dose of GNR. Moreover, this was achieved on a tumor model with relevant characteristics for the clinical translation prospects. As opposed to the subcutaneous tumors of the majority of PPTT published studies, orthotopic implantation provides a better environment for the expression of tumorigenicity of the cells, higher blood perfusion, larger vessel diameters and lower interstitial fluid pressure (IFP) [22, 121]. PPTT has been tested in only a handful of orthotopic models before, syngeneic in mice and rats [122–124] or human-derived [125]. Furthermore, ccRCC is of clinical relevance due to the limited treatment strategies available today for this type of tumor. It is generally resistant to chemotherapy and radiotherapy, so the first-line treatments have been achieved more recently, with the development of antiangiogenic therapy and its combination with immunotherapy [49, 126, 127]. Still, resistance to them is developed over time reducing their long-term efficacy while other patients are not eligible for them.

In addition, our experiments were conducted on tumors of dimensions larger than the typical optical penetration depth and heat diffusion characteristic length, and have thus allowed us to observed the effect of partial treatment of tumor volumes and relate it to the physical mechanisms causing it. On the counterpart, a 1 cm^3^ tumor represented a much higher relative burden for a mouse than it would in a larger animal or a human, which may affect the well-being, stability, response to therapy and survival of the individuals.

Further capabilities of the paradigm we have presented could be achieved with tumor diffuse optical measurements during the irradiation and hyperthermia, through non-contact implementations like those in [128, 129]. Following the alterations in these parameters without disrupting the therapy would be a powerful tool for the fine adjustment of the light dose in real time, as has been done in PDT experiments by reacting to the changes in tumor *BFI* [41].

## Conclusion

Better-informed PPTT is required to understand the variability in outcome and the mechanisms behind it, in order to build optimized and reliable treatment protocols. Our methods have successfully characterized the outcome of the therapy with standard techniques, identified the relevance of optical parameters, NP concentration and water fraction for therapy response and provided validated extrapolated markers of therapy outcome through simplistic simulations. In principle, the same methods could be used to study any other set of therapy conditions and protocols that cannot be simulated, such as different tumor types, nanoparticle formulations, number of irradiation courses or combinations with other treatment methods. A training cohort of xenografted mice would allow to establish the modeling of therapy, like shown here, coarse-tune the therapy parameters and develop personalization protocols. This work thus shows the capabilities of our methodology to optimize PPTT and accelerate its progress into clinical practice.

## Supporting Information

A supplemental document is available online with the preprint at bioRxiv repository.

## Acknowledgements

This project has received funding from Fundació CELLEX Barcelona, Fundació Mir-Puig, Agencia Estatal de Investigación, the Secretaria d’Universitats i Recerca del Departament d’Empresa i Coneixement de la Generalitat de Catalunya, Secretaria d’Universitats i Recerca del Departament de Recerca i Universitats de la Generalitat de Catalunya (AGAUR-2019-FI B-00968), the European Social Fund (L’FSE inverteix en el teu futur)—FEDER and Fundació Joan Ribas Araquistain (FJRA).

