## Supplementary information for "A Toolbox for the Personalization of Plasmonic Photothermal Therapy in Orthotopic Mouse Tumor Models"

May 2, 2025

#### 1 GNR synthesis details

Gold nanorods (GNR) were grown in-house using a seed-mediated method in cetyltrimethylammonium bromide (CTAB, Sigma-Aldrich) as described in detail in reference [1]. The GNR suspension in CTAB was ultra-centrifuged (Avanti J-E rotor JA-25.50, Beckman Coulter, USA) for 25 min at 23708 xg (14000 rpm) and mantained at 31°C; then the excess of CTAB was removed by eliminating 90% of the supernatant. Next, the ligand exchange reaction was carried out by mixing in double purified water (MilliQ, Merck KGaA, Germany) 50% v/v of citrate 0.1 M pH 4 (C3434, Sigma-Aldrich, USA), 3000 Da polyethylene glycol (PEG, PEG1099, Iris Biotech, Germany) at 1 mg/mL and the concentrated GNRs-CTAB at 0.25 OD/mL. The mixture was sonicated for 30 min and kept at 28°C overnight to allow the exchange of CTAB chains and PEG moieties. The PEGylated solution (GNR–PEG) was ultra-centrifuged (25 min at 23708 xg and 28°C) to remove unbounded PEG moieties, filtered (25 mm regenerated cellulose syringe filter, 0.2 µm pore size, Corning Inc, USA) to discard aggregation, washed twice and diluted with ultrapure water (Millipore, Merck, Germany) to the desired final concentration. The final size of the GNRs was 11 nm x 44 nm.

The absorption spectra of GNR–PEG in suspension were measured prior to injection using a well-plate spectrophotometer (Synergy H1, BioTek Instruments, now Agilent, USA) with 96 well plates (polystyrene, clear, flat bottom, Nunc MicroWell, Thermo Fisher Scientific) with 0.69 cm path length (250 µl of studied volume). The longitudinal surface plasmon resonance of the GNRs was located around 810 nm, as determined by their size and aspect ratio. Water spectrum subtraction and triplicates were done for each sample. GNR concentration was determined as $C = Abs/(L\epsilon)$, where $C$ $(mol/l)$ is the molar concentration, $Abs$ (cm$^{-1}$) is the optical absorbance (natural logarithm), $L$ (cm) is the path length of the cuvette and $\epsilon$ $(cm^{-1}(mol/l)^{-1})$ is the GNR molar extinction coefficient.

#### 2 Analytes for chronic toxicity study

Twenty-two parameters were selected and analyzed from blood samples. Their values for mice injected with 21 nM GNR–PEG and control mice injected with PBS are presented in table S1. Wilcoxon rank-sum tests were used for differences between groups. There were no differences between groups considering Bonferroni correction.

Table S1: Blood sample analysis results for long-term GNR toxicity study. Values expressed as mean ± SD. Uncorrected $p$-values are shown.

| **Parameter** | $N_{GNR}$ | $N_{PBS}$ | **GNR–PEG** | **PBS** | $p$**-value** |
|---|---|---|---|---|---|
| Alanine aminotransferase (U/l) | 8 | 6 | 81.2 ± 89.3 | 56.6 ± 29.6 | 0.66 |
| Aspartate aminotransferase (U/l) | 8 | 6 | 378 ± 508 | 210 ± 150 | 0.75 |
| Creating kinase (U/l) | 7 | 6 | 2528 ± 3013 | 1465 ± 1518 | 0.95 |
| Creatinine (mg/dl) | 8 | 6 | 0.262 ± 0.038 | 0.26 ± 0.02 | 0.89 |
| Fe (µg/dl) | 8 | 6 | 175.7 ± 74.8 | 232.8 ± 25.5 | 0.06 |
| Alkaline phosphatase (U/l) | 8 | 6 | 21.5 ± 16.9 | 44.4 ± 18.7 | 0.08 |
| Glucose (mg/dl) | 8 | 6 | 263 ± 182 | 293.6 ± 96.5 | 0.23 |
| K (mmol/l) | 8 | 6 | 7.11 ± 1.07 | 6.11 ± 1.15 | 0.14 |
| Proteins (g/dl) | 8 | 6 | 5.29 ± 0.33 | 5.63 ± 0.63 | 0.23 |
| Sodium (mmol/l) | 8 | 6 | 152.55 ± 2.41 | 151.35 ± 3.13 | 0.48 |
| White blood cells, peroxidase method ($10^3$ cell/µl) | 5 | 4 | 3.0 ± 2.5 | 4.71 ± 1.45 | 0.41 |
| White blood cells, basophil method ($10^3$ cell/µl) | 5 | 4 | 3.16 ± 1.96 | 5.16 ± 1.71 | 0.19 |
| Red blood cell count ($10^6$ cell/µl) | 5 | 4 | 8.99 ± 1.69 | 9.39 ± 1.21 | 0.73 |
| Hematocrit (%) | 5 | 4 | 38.54 ± 7.48 | 45.63 ± 4.51 | 0.18 |
| Platelets ($10^3$ cell/µl) | 5 | 4 | 1008 ± 660 | 1322 ± 47 | 0.73 |
| Neutrophils (%) | 5 | 4 | 36.6 ± 14.0 | 21.0 ± 4.5 | 0.11 |
| Lymphocytes (%) | 5 | 4 | 39.7 ± 19.8 | 68.0 ± 7.1 | 0.11 |
| Monocytes (%) | 5 | 4 | 2.56 ± 1.36 | 3.6 ± 1.57 | 0.46 |
| Eosinophils (%) | 5 | 4 | 19.34 ± 14.56 | 6.5 ± 3.88 | 0.03 |
| Large unstained cells (%) | 5 | 4 | 1.64 ± 2.11 | 0.70 ± 0.41 | 0.54 |
| Basophils (%) | 5 | 4 | 0.30 ± 0.34 | 0.175 ± 0.096 | 0.9 |

### 3 Diffuse reflectance spectroscopy analysis

#### 3.1 Diffuse reflectance model

Broadband DRS data was fitted using the delta-Eddington $P_1$ approximation ($\delta$-$P_1$) to the radiative transport equation for a homogeneous medium and a pencil source, as detailed in reference [2]. This model is able to describe light reflectance in the range of SDSs of 0.5 – 5 mm better than the diffusion approximation [3].

Under this model, the macroscopic behavior of light is determined by the values of the absorption coefficient $\mu_a$, the scattering coefficient $\mu_s$ and two scaling relations, namely, the usual definition for the reduced scattering coefficient $\mu_s' = (1-g)\mu_s$ and the additional $\mu_s^* = \gamma\mu_s'$, with both factors $g$ $(0 < g < 1)$ and $\gamma$ $(1 < \gamma < 2)$ dependent on the anisotropy of the scattering phase function. The coefficient $\mu_s^*$ plays the role of $\mu_s$ for the non-forward-scattered light, leading to an analogue inhomogeneous diffusion equation. Using its Green's function and extrapolated boundary conditions, the solutions for the diffused fluence rate $\Phi_\mathrm{d}(\rho, z)$ take the form:

$$\Phi_\mathrm{d}(\rho, z) = \frac{v}{4\pi D}\frac{\mu_\mathrm{s}^*}{(\mu_a + \mu_\mathrm{s}^*)}\int_0^\infty \left[\frac{e^{-r_1/L_D}}{r_1} - \frac{e^{-r_2/L_D}}{r_2}\right] e^{-(\mu_a+\mu_\mathrm{s}^*)z'}\mathrm{d}z', \tag{1}$$

where $\rho$ is the source-detector separation, $z$ is the inward spatial dimension into the tissue, $v$ is the speed of light in the medium, $D = v/3(\mu_s' + \mu_a)$ is the light diffusion coefficient, $L_D = (3\mu_a(\mu_s' + \mu_a))^{-1/2}$ is the optical penetration depth and the distances to the source points at $z = z'$ are $r_1 = \sqrt{(z-z')^2 + \rho^2}$ and $r_2 = \sqrt{(z - z' - 2z_b)^2 + \rho^2}$ (with $z_b \sim 6.51D$). The solution for the diffuse reflectance $R(\rho)$ then becomes [2]:

$$R(\rho) = -\frac{D}{vS_0}\left.\frac{\partial\Phi_d(\rho, z)}{\partial z}\right|_{z=0} = \frac{\Phi_d(\rho, z = 0)}{2A}. \tag{2}$$

##### 3.2 Fitting algorithm

DRS data was fitted following the self-calibration method used in references [4, 5] and implementing an iterative fitting algorithm. Prior to fitting, the signals were dark-subtracted and normalized using the data from other source-detector pairs, as follows. First, the dark-subtracted signal of each collector, $I_j(\lambda)$ $(j = 1, ..., 6)$, was divided by the calibration measurement $I_j^{\mathrm{cal}}(\lambda)$ (same collection fiber, illumination through calibration source) in order to remove collector–skin coupling effects and the source spectrum factor. Later, this ratio was normalized to that of the previous collector in order to minimize source–skin coupling effects [6]. This can be summarized for collectors $j = 2, ..., 6$ as:

$$I_j^{\mathrm{fit}}(\lambda) = \frac{I_j(\lambda)/I_j^{\mathrm{cal}}(\lambda)}{I_{j-1}(\lambda)/I_{j-1}^{\mathrm{cal}}(\lambda)} \tag{3}$$

The five resulting normalized signals $I_j^{\mathrm{fit}}(\lambda)$ were fitted separately (assuming a homogeneous medium each time), as the probed volume is different for each pair and the tissue properties can be heterogeneous. The model fitted to $I_j^{\mathrm{fit}}(\lambda)$ was thus $R(\rho_j)/R(\rho_{j-1}) = \Phi_{\mathrm{d}}(\rho_j, 0)/\Phi_{\mathrm{d}}(\rho_{j-1}, 0)$.

Effective Mie scattering was assumed for the spectral dependence of $\mu_s'$, of the form $\mu_s'(\lambda) = A(\lambda/\lambda_0)^{-b}$, where the scattering amplitude $A$ and the scattering power $b$ were fitted experimentally and the reference wavelength $\lambda_0 = 785$ nm was used [7, 8].

For the absorption coefficient $\mu_a$, linear contributions were considered from the main chromophores, Hb, $HbO_2$, $H_2O$ and GNRs, using $\mu_a(\lambda) = \sum_i C_i \epsilon_i(\lambda)$, where $\epsilon_i(\lambda)$ is the absorption coefficient per unit molar concentration for chromophore $i$ and $C_i$, its molar concentration. As an exception, water contribution is expressed as $\mu_{a,H_2O} \times c_{H_2O}$ where $c_{H_2O}$ is the water fraction (dimensionless), and $\mu_{a,H_2O}$, the absorption coefficient of pure water. The total hemoglobin concentration $THC = C_{HbO_2} + C_{Hb}$ and tissue oxygen saturation $StO_2 = C_{HbO_2}/THC$ were fitted parameters together with $C_{GNR}$ and $c_{H_2O}$. The full set of DRS fitted parameters was thus $\{A$, $b$, $c_{H_2O}$, $THC$, $StO_2$, $C_{GNR}\}$.

The fitting of the model was validated on liquid phantoms and an algorithm for objectively assessing the fit quality of the in vivo data was tested and optimized through the analysis of residuals.

Different wavelength bands were used for spectral fitting to reduce the crosstalk between absorption and scattering. Preestablished or data-based wavelength selection have been proposed before based on available wavelengths, optical properties, chromophores considered and levels of noise [9–13]. In this study a fixed set of wavelength bands was used, given that they performed well in tumor and shoulder data, with or without GNR presence.

To reduce the effects of possible local minima, an iterative approach was chosen in which each variable was fitted roughly in the wavelength range were they had a higher impact on the reflectance, while leaving the rest of parameters constant. In addition, the first derivative of the signal with respect to wavelength was exploited to improve the determination of $c_{H_2O}$ around the absorption peak at 970 nm, as this helped to decouple it from $C_{GNR}$ by enhancing its spectral features and further reducing any possible skin coupling effects [6, 14, 15]. The derivative was calculated using smoothed signals with a Savitzky-Golay filter (2nd degree polynomial, 50 nm window).

As the first step in the algorithm, all six parameters were fitted on the normalized signal $I_j^{\mathrm{fit}}(\lambda)$ in the range of $635 - 1000$ nm as a starting point. The next steps were as follows:

**II.** fit for $\{StO_2, THC\}$ using $I_j^{\mathrm{fit}}(\lambda)$ (0th derivative) in the $690 - 850$ nm range,

**III.** fit for $\{A, b, C_{GNR}\}$ using $I_j^{\mathrm{fit}}(\lambda)$ in the $650 - 850$ nm range, and

**IV.** fit for $c_{H_2O}$ using the 1st derivative of $I_j^{\mathrm{fit}}(\lambda)$ in the $930 - 1000$ nm range.

The listed steps were reiterated until the $\chi^2$ goodness-of-fit statistic reduced by less than 5% in a cycle (II – IV) compared to the previous one (limited to 255 cycles to discard possible non--convergence). In each step, a non-linear least-squares trust-region-reflective algorithm was used in MATLAB software (Version 9.5.0.1067069 (R2018b) Update 4, The Mathworks, Inc.). This resulted in relative residuals with respect to $I_j^{\text{fit}}(\lambda)$ ($< 1\%$). Fittings where they surpassed the 1% threshold at one or more wavelength values (in 0th-derivative steps) were discarded (3 – 8 % of fittings, depending on condition).

### 4 Other statistical details

#### 4.1 Group tests

The pair-wise comparisons between treated groups in final skin temperature, histological results and diffuse optical parameters were done using non-parametric Wilcoxon rank-sum tests. Paired data of gold content in different organs of mice was tested with Wilcoxon signed-rank test.

#### 4.2 Correlations with skin temperature

Linear regressions between $T_{ROI}$ and other variables were done using the Deming regression method. For optical data, the median of the values was used. The angle for the least-squares minimization was calculated as the ratio of the average standard deviations for each variable.

#### 4.3 GNR accumulation model

To relate optical data before and after injection, linear mixed-effects (LME) models were used to account for the repeated measures of optical data on the tumor and for the different mice in each group. The mouse ID was used as a random effect nesting data from repeated probe replacements. As there was no significant effect of the SDS, the data of different detectors was considered as repeated measures as well.

Significance of the covariates was evaluated through likelihood ratio tests with respect to the corresponding null model lacking such effect (significance level of $p < 0.05$). Residual normality was not a requisite as it has a negligible effect on the estimates [16, 17].

#### 4.4 Correction for multiple comparisons

All multiple comparison tests were weighted with conservative Bonferroni correction, i.e. corrected $p$-values were considered $p = m \cdot p'$, where $m$ is the number of pairwise tests done on the same data and $p'$ is the usual $p$-value. Significance was evaluated as $p < 0.05$.

### 5 Light transport and heat diffusion simulation parameters

Table S2 summarizes all the parameters used for light transport and heat diffusion simulations through MC Matlab. They were considered to first approximation as constant under temperature change.

Table S2: Parameters used in light transport and heat diffusion simulations.

| **Simulation parameters** | |
|---|---|
| Voxels | 101 × 101 × 101 |
| Total dimensions | 5 × 5 × 5 cm |
| MC iterations | $10^7$ |
| MC source beam diameter | 1.2 cm |
| MC $\lambda_T$ | 808 nm |
| Heat diffusion total duration | 1000 s |
| Heat diffusion time step | 0.5 s |

| **Tissue properties** | |
|---|---|
| Refraction index | 1.4 |
| $\mu_a(\lambda_T)$ | Mouse data |
| $\mu_s(\lambda_T)$ | Mouse data |
| g | 0.9 |
| Mass specific heat | 4.2 J/(K × kg) |
| Mass density | 1.1 kg/cm$^3$ |
| Thermal conductivity | 5.5 $10^{-3}$ W/(K × cm) |

| **Values sampled for extrapolation of therapy conditions** | |
|---|---|
| $\mu_a(\lambda_T)$ | 0.1 – 0.9 cm$^{-1}$ |
| $\mu_s(\lambda_T)$ | 80 cm$^{-1}$ |
| Treatment power | 0.5 – 3.5 W |
| Treatment duration | 100 – 600 s |
| Total delivered energy | 50 – 2100 J |

### 6 Individual tumor growth data

Tumor volume evolution after therapy was fitted with an exponential model (LME) to look for group differences, as described in Methods section of the manuscript, with a random slope for each mouse. The fitted models using estimates for individual mice are shown in **Figure S1** for the three experimental groups (controls, PBS and GNR) to display variability in tumor evolution between groups.

### 7 H&E staining results

Hematoxylin and Eosin (H&E) staining of excised tumor samples was conducted to quantify the necrosis present in tumor cells for mice in the three experimental groups (controls, PBS and GNR) in the short-term (1 day) and long-term (15 – 17 days) after PPTT. Results are shown in **Figure S2**. No significant differences were found between groups in the short- or long-term. No significant differences were found for any group between short and long terms.

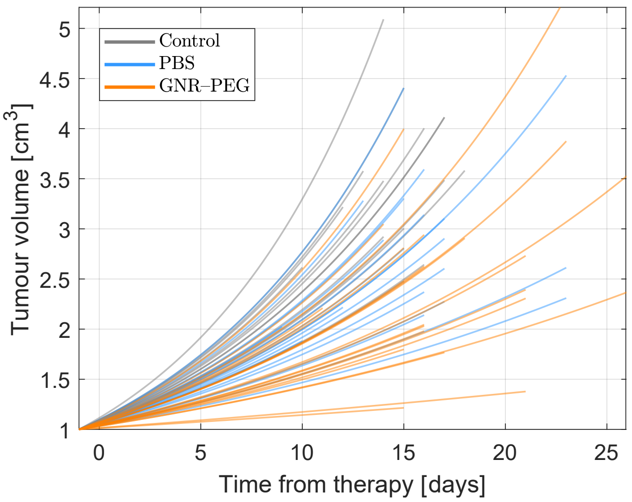


Figure S1: Fitted LME model tumor volume curves (long-term cohorts: $N_{\text{control}}$=15, $N_{\text{PBS}}$=19, $N_{\text{GNR}}$=21). Each curve shows the fitted values for a mouse, plotted up to maximum survival day.

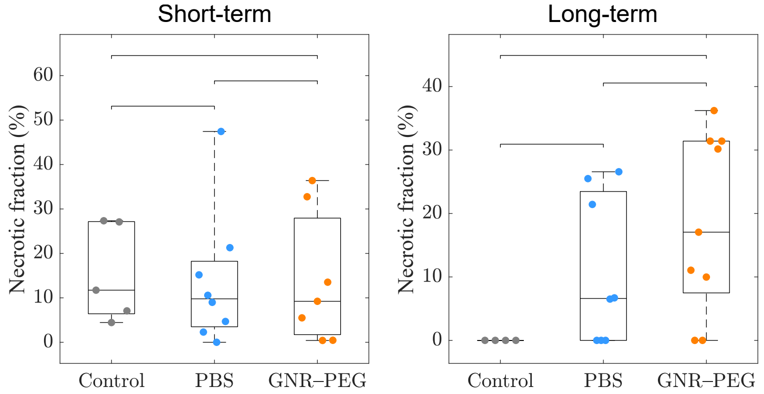


Figure S2: Tumor Necrotic fraction for mice sacrificed at a fixed timepoint of 1 day after therapy (short term, $N_{\text{control}}$=4, $N_{\text{PBS}}$=8, $N_{\text{GNR}}$=9) and 15 – 17 days after PPTT (long term, $N_{\text{control}}$=5, $N_{\text{PBS}}$=8, $N_{\text{GNR}}$=7). The boxes indicate the medians and interquartile ranges, the whiskers show the full ranges and the brackets indicate the pairwise tests conducted (Wilcoxon rank-sum test with Bonferroni correction). No statistical differences were found in these tests.

#### 8 Correlation between maximum skin temperature and tumor total hemoglobin concentration

Figure S3 shows the correlation between $T_{ROI}$ and $THC$ for each treated group ($p < 10^{-3}$). Deming regression revealed for the GNR–PEG group an intercept of 50 ± 2°C and a slope of 0.05 ± 0.02°C/µM and, for the PBS group, an intercept of 40 ± 2°C and a slope of 0.10 ± 0.03°C/µM.

#### 9 Relation between mouse survival after therapy and body weight

Mouse survival was associated to mouse body weight at the day of therapy (24.7 – 41.3 g, $p = 0.006$) in addition to the experimental group (control, PBS or GNR–PEG). The fitted estimates for the log(HR) were $-0.6 \pm 0.5$ for the PBS group, $-1.3 \pm 0.5$ for the GNR–PEG group, both

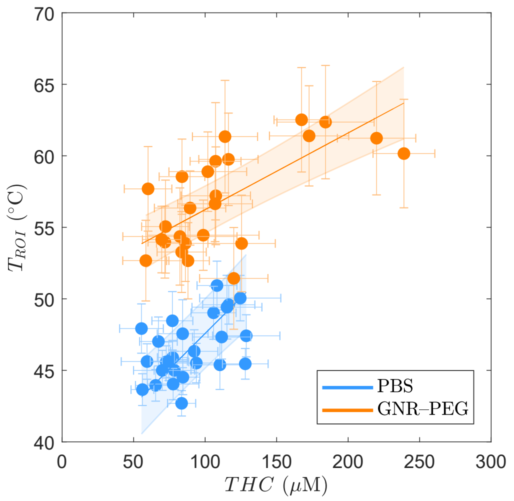


Figure S3: Relationship between $T_{ROI}$ values (IR thermography) and pre-therapy tumor $THC$ (DRS), overlayed with the Deming regression model for each group ($n_{\mathrm{GNR}}$=30, $n_{\mathrm{PBS}}$=27)). Each point is the median value for a mouse with standard deviation bars. The shaded regions indicate the SE range.

with respect to controls, and $-0.18 \pm 0.07$/g for the slope with weight. **Figure SS4** shows the probabilistic index for the fitted Cox model.

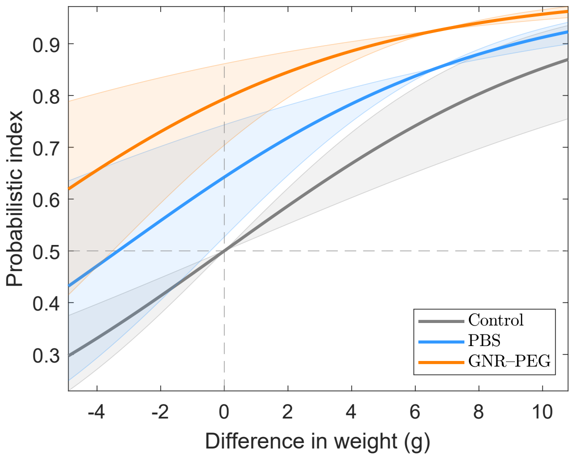


Figure S4: Probabilistic index for the Cox model relating mouse survival with body weight and experimental group, relative to the control group. The range plotted is the observed range of absolute differences between subjects (long-term cohorts: $N_{\mathrm{control}}$=15, $N_{\mathrm{PBS}}$=19, $N_{\mathrm{GNR}}$=21). The shaded regions indicate $e^{-1}$ confidence intervals.

### 10 Changes in hemodynamic parameters after therapy

**Figure S5** shows the changes in hemodynamic parameters for the three experimental groups, denoted as "$\Delta x$" for parameter "$x$". In tumor tissue, the changes in all variables were different for the GNR–PEG group to those of controls ($p < 0.02$), while the PBS group only showed a difference in $\Delta StO_2$ with respect to controls ($p = 0.04$). Between the treated groups, there were significantly different variations in $c_{H_2O}$ ($p < 10^{-3}$), $THC$ ($p = 0.01$) and $BFI$ ($p = 0.01$). All of the changes were greater in magnitude for the GNR–PEG group.

On the other hand, in the shoulder there were significant differences for each treated group with respect to controls in $\Delta StO_2$ ($p < 0.04$) and $\Delta BFI$ ($p < 3 \cdot 10^{-3}$).

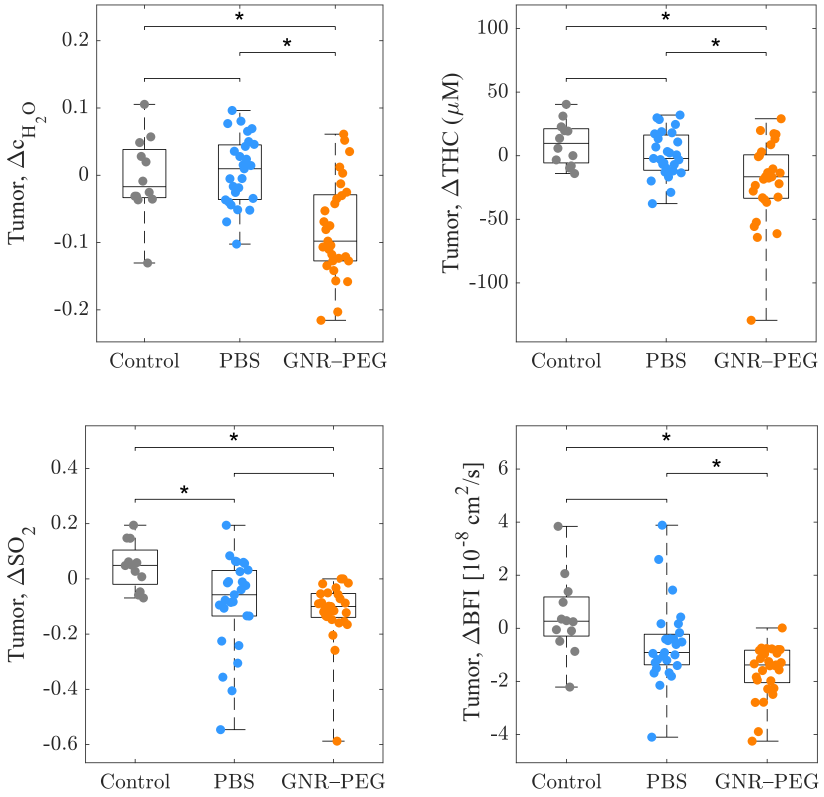


Figure S5: Variation in tumor hemodynamic parameters from pre- to post-therapy for the three experimental groups ($N_{\text{control}}$=19, $N_{\text{PBS}}$=27, $N_{\text{GNR}}$=30). The boxes indicate the medians and interquartile ranges, the whiskers show the full ranges and the brackets indicate the pairwise tests conducted (Wilcoxon rank-sum test with Bonferroni correction, *: $p$<0.05).

#### 11 Relation between optical penetration depth, skin temperature and treated volume

Experimental evidence showed that a lower optical penetration depth ($L_D = [3\mu_a(\mu_s' + \mu_a)]^{-1/2}$) measured before therapy was related to higher skin temperature values, despite implying that the treatment light will visit a smaller, shallower volume of tissues during irradiation. This contradictory behavior was studied with Monte Carlo light transport and heat diffusion simulations. A set of typical absorption coefficient values ($\mu_a$=0.2 – 0.8 cm$^{-1}$) and reduced scattering coefficient values ($\mu_s'$=7 – 10 cm$^{-1}$) were chosen to study the relation of $L_D$ with simulated variables describing the heating: $T_{ROI}^{sim}$ and $V_{43}^{sim}$. The therapy used were those of the experiments: 2 $W/cm^2$ intensity, 12 $cm$ diameter area and 300 $s$ irradiation.

Simulations showed that for both, $T_{ROI}^{sim}$ and $V_{43}^{sim}$ there is a dominant effect of $\mu_a$ for the range of values simulated. Higher $\mu_a$ increases their values despite reducing $L_D$, as observed experimentally. This can be understood through the fact that the total amount of heat generated

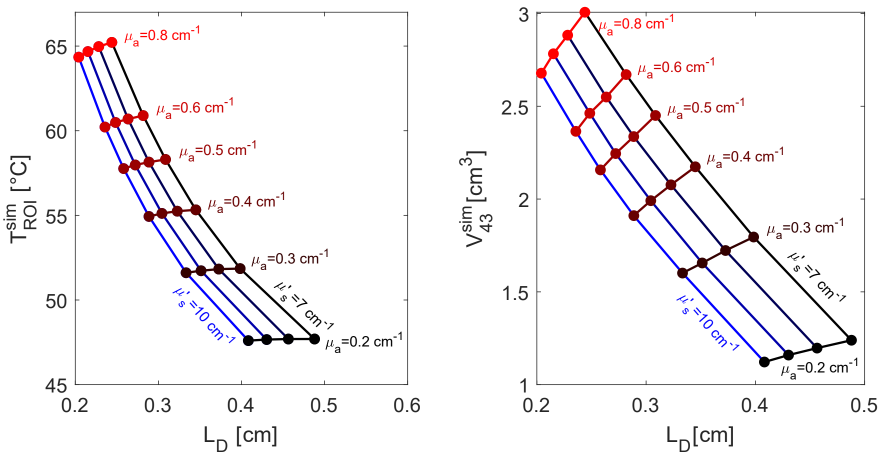


Figure S6: Dependence of simulated $T_{ROI}^{sim}$ and $V_{43}^{sim}$ on $L_D(\lambda_T)$, for $\mu_a(\lambda_T)$ and $\mu_s'(\lambda_T)$ values sampled over the experimental ranges.

is greatly dependent on $\mu_a$, and the energy will reach the skin and deeper volume of the tumor through heat conduction, and not light penetration. Note that values of $\mu_a$ between 0.3 and 0.8 $cm^{-1}$ represent tumor tissue with accumulated GNRs, supporting the mechanism of absorption enhancement for better treatment efficacy.

This is opposite to the contribution of $\mu_s'$, which shows the intuitive result of lower $V_{43}^{sim}$ for a lower light penetration due to higher scattering (for a fixed absorption value). Moreover, the same is true for $T_{ROI}^{sim}$. This can be explained by the fact that heat generation is not only dependent on $\mu_a$ values, but also on the fluence rate distribution on the tissue. A higher scattering will mean more light will escape from the tissue and thus, it will not be absorbed and converted into heat.

#### 12 Extrapolation of absorption values

**Figure S7** shows extrapolation of simulated parameters $T_{ROI}^{sim}$, $V_{43}^{sim}$ and $V_{55}^{sim}$ to absorption values in the 0.2 – 0.8 cm$^{-1}$ range, irradiation powers of 0.5 – 3 W and irradiation time of 300 s. These results together with irradiation time and extrapolation (Figure 5 on main document) can be used for therapy planning.

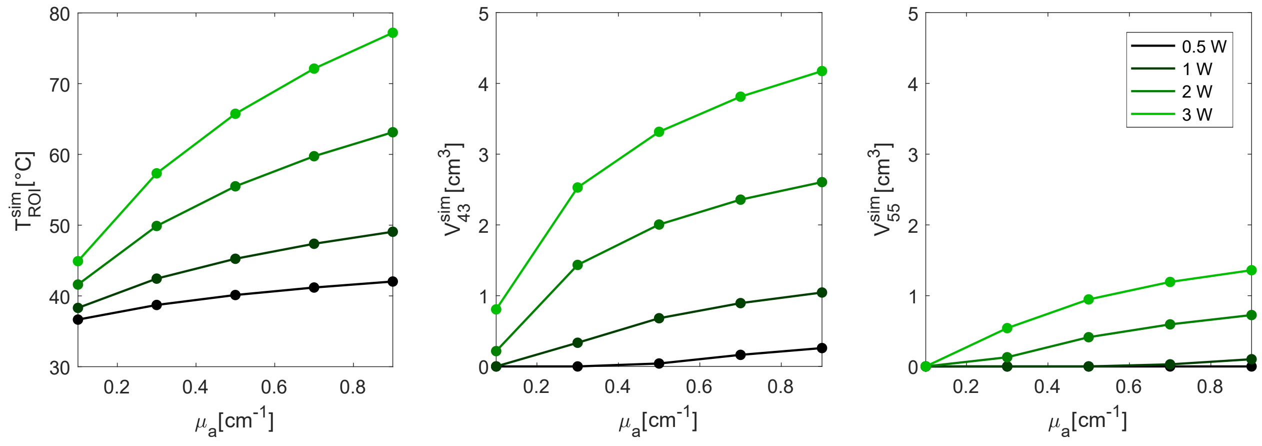


Figure S7: Dependence of simulated $T^{sim}_{ROI}$, $V^{sim}_{43}$ and $V^{sim}_{55}$ on extrapolated $\mu_a(\lambda_T)$ values for different incident powers ($\mu'_s(\lambda_T)$=10 cm$^{-1}$, 300 s irradiation).
